# *Arabidopsis thaliana* BBX14 is a target of GLK1 and involved in high-light acclimation, photomorphogenesis and GUN-type retrograde signaling

**DOI:** 10.1101/2023.03.03.530939

**Authors:** Vasil Atanasov, Julia Schumacher, Jose M Muiño, Catharina Larasati, Liangsheng Wang, Kerstin Kaufmann, Dario Leister, Tatjana Kleine

**Author notes:** Corresponding author: Tatjana Kleine.

## Abstract

Development of photosynthetically competent seedlings requires both light and retrograde biogenic signaling pathways. The transcription factor GLK1 functions at the interface between these pathways, and receives input from the biogenic-signaling integrator GUN1. BBX14 was previously identified, together with GLK1, in a core module that mediates the response to high light levels and biogenic signaling. To gain insight into the function of BBX14, we generated *BBX14* overexpressors and CRISPR/Cas-mediated *bbx14* mutant plants, conducted high-light, RT-qPCR and ChIP-Seq experiments, measured photosynthetic parameters, chlorophyll contents and growth rates, and analyzed alterations in transcriptomics. We found that, although overexpression of BBX14 is deleterious under normal growth conditions, BBX14 is needed to acclimate plants to high light stress. *BBX14* is a direct target of GLK1, and RNA-Seq analysis suggests that BBX14 is involved in the circadian clock. Knockout of *BBX14* results in a long-hypocotyl phenotype that depends on a retrograde signal, and *BBX14* expression during biogenic signaling requires GUN1. Finally, we clarify the role of BBX14 in GUN-type biogenic signaling. We conclude that BBX14 is an integrator of photomorphogenetic and biogenic signals, and suggest that BBX14 is a nuclear target of retrograde signals downstream of the GUN1/GLK1 module.

## INTRODUCTION

Light is indispensable for plants. Apart from driving photosynthesis, it is required to direct growth, as well as developmental and acclimation processes. For example, in darkness, etiolated/skotomorphogenic Arabidopsis seedlings develop long hypocotyls, unexpanded and appressed cotyledons with etioplasts, and an apical hook that protects the apical meristem from damage (Han *et al*., 2020). On the other hand, photomorphogenetically de-etiolated seedlings change their form to maximize light capture for photosynthesis by inhibiting hypocotyl elongation, unfolding the hook, stimulating cotyledon separation and expansion, and activating the formation of fully functional chloroplasts (Von Arnim & Deng, 1996). At the heart of the light-dependent signal transduction network that controls this transition is the E3 ubiquitin ligase CONSTITUTIVE PHOTOMORPHOGENIC 1 (COP1). COP1 integrates signals from photoreceptors and regulates a set of downstream components including the transcription factors ELONGATED HYPOCOTYL 5 (HY5) (Podolec & Ulm, 2018; Han *et al*., 2020), GOLDEN2-LIKE 1 (GLK1) and GLK2, which regulate genes involved in chlorophyll biosynthesis and formation of the photosynthetic apparatus (Waters *et al*., 2009), and B-box (BBX) proteins (Xu, 2020).

In plants, BBX proteins form a subgroup of zinc-finger transcription factors (Talar & Kielbowicz-Matuk, 2021), which is comprised of 32 members in *Arabidopsis thaliana* (Khanna *et al*., 2009). The B-box itself can mediate a variety of functions including protein-protein interactions (Datta *et al*., 2006) and activation of transcription (Datta *et al*., 2007). The first BBX protein identified in Arabidopsis was CONSTANS (CO, BBX1), which promotes flowering (Putterill *et al*., 1995). In subsequent studies, very different names were used for members of the B-box family, which prompted Khanna et al. (2009) to propose a uniform nomenclature for them. These authors numbered them from BBX1 to BBX32, and sorted them into clades I to V, based on multiple sequence alignment, phylogenetic analysis and the presence of additional domains. Clades I (BBX1-BBX6) and II (BBX7-BBX13) are characterized by two B-boxes and a CCT (CO, CO-LIKE and TOC1) domain, and clade III (BBX14-BBX17) by a single B-box and a CCT domain, while members of clade IV (BBX18-BBX25) have two B-boxes, and those of clade V (BBX26-BBX32) have one. It is now known that BBX proteins participate in the regulation of plant growth and development, hormonal pathways, and biotic and abiotic stress tolerance (Khanna *et al*., 2009; Gangappa & Botto, 2014; Vaishak *et al*., 2019; Xu, 2020; Talar & Kielbowicz-Matuk, 2021). Moreover, different members of the group can act antagonistically, serving as positive and negative regulators of the same processes. For example, during seedling photomorphogenesis, BBX4 and BBX20-BBX23 (Datta *et al*., 2006; Datta *et al*., 2007; Datta *et al*., 2008; Fan *et al*., 2012; Zhang *et al*., 2017) have been identified as positive, and BBX18, BBX19, and BBX24, BBX25, and BBX28-BBX32 (Khanna *et al*., 2006; Kumagai *et al*., 2008; Holtan *et al*., 2011; Heng *et al*., 2019; Cao *et al*., 2022) as negative regulators, respectively.

The development of photosynthetically competent seedlings is strictly dependent on the exchange of information between plastids and the nucleus. Since most chloroplast proteins are encoded in the nucleus, it exercises anterograde control over the plastids (Jan *et al*., 2022). Conversely, during retrograde signaling, chloroplasts emit signals that convey information relating to their status to the nucleus, so that nuclear gene expression can be adjusted accordingly (Kleine & Leister, 2016; Jan *et al*., 2022; Liebers *et al*., 2022). This mechanism is exemplified by the response of seedlings to treatment with norflurazon (NF, an inhibitor of carotenoid biosynthesis) or inhibitors of organellar protein synthesis such as lincomycin (LIN): each of these agents reduces levels of transcripts derived from so-called photosynthesis-associated nuclear genes (*PhANG*s) (Oelmuller *et al*., 1986). Although the quest for *genomes uncoupled* (*gun*) mutants (i.e. seedlings that continue to express *PhANG*s even though plastid development has been blocked by NF or LIN) began 30 years ago, the mechanisms underlying this phenotype have yet to be fully clarified, and the number of known *gun* mutants now stands at less than ten (Richter *et al*., 2023).

BBX14 (COL6), a member of clade III, attracted our attention because it was found together with GLK1 and GLK2 in a core module required for the nuclear retrograde response to altered organellar gene expression (Leister & Kleine, 2016). The identification of this module was based on database analyses, which indicated that changes in gene expression comparable to those provoked by treatment with LIN, NF or high light (HL) also take place in two mutants that are defective in plastid gene expression. Using this module, overexpressors of GLK1 and GLK2 were shown to be *gun* mutants under both NF and LIN conditions (Leister & Kleine, 2016). Moreover, based on a co-expression network, BBX14 was suggested as a potentially important regulator of the response to high light (HL) levels (Huang *et al*., 2019).

The present study was designed to gain further insight into the function of BBX14. Our new data show that overexpression of BBX14 is deleterious under normal growth conditions, while *bbx14* plants behave like wild type (WT). However, lack of BBX14 compromises acclimation of plants to HL. We identified *BBX14* as a direct target of GLK1, and RNA-Seq analysis of *bbx14* mutant seedlings reveals specific de-regulation of transcripts encoding proteins involved in the circadian clock. BBX14 promotes photomorphogenesis, and the long hypocotyl and root phenotype seen in *bbx14* is comparable to that of one of the *glk1* mutant. This phenotype in turn depends on a retrograde signal, and reduction of *BBX14* mRNA levels during biogenic signaling depends on GUN1. Transcript levels of several, but not all of the *PhANG* genes investigated are de-repressed under NF treatment, suggesting that BBX14 is a candidate for the reception of retrograde signals downstream of GLK1.

## MATERIALS AND METHODS

### Plant material and growth conditions

All the lines used were in the Col-0 background, and the mutants *bbx14-1* (SAIL_1221_D02; N878600) and *pifq* (N2107737) were obtained from the NASC and ABRC, respectively. The T-DNA insertion in *bbx14-1* was confirmed with the primers listed in Supporting Information Table S1 (see also Fig. S1). The mutants *cry1* (Kleine *et al*., 2007), *gun1-102* (Tadini et al., 2016), *gun4-1* (Susek et al., 1993), *glk1* and *glk2* (Waters *et al*., 2009) mutants have been described previously. The inducible overexpression line (Coego *et al*., 2014) was obtained from the ABRC stock center.

Growth conditions employed in the experiments are described in detail in Supporting Information Methods S1.

### Nucleic acid extraction

For DNA isolation, leaf tissue was homogenized in extraction buffer containing 200 mM Tris/HCl, pH 7.5, 25 mM NaCl, 20 mM EDTA and 0.5% (w/v) SDS. After centrifugation, DNA was precipitated from the supernatant by adding isopropanol. After washing with 70% (v/v) ethanol, the DNA was dissolved in distilled water.

For RNA isolation, frozen tissue was ground in liquid nitrogen. Total RNA was extracted using Direct-zol™ RNA MiniPrep Plus columns (Zymo Research, Irvine, USA) following the manufacturer’s instructions. RNA quality and concentration and the A_260_/A_280_ ratio were assessed by agarose gel electrophoresis and spectrophotometry. Isolated RNA was stored at −80°C prior to use.

### cDNA synthesis and quantitative real-time PCR analysis

First-strand cDNA synthesis and RT-qPCR were performed essentially as described previously (Wang *et al*., 2022), except that PCRs were carried out in the CFX Connect real-time system (Bio-Rad, Munich, Germany).

### Generation of *BBX14* CRISPR and overexpression lines

The pHEE401-E vector, which provides an egg-cell-specific promoter, was used to generate the *CRISPR-Cas* lines *bbx14-2* and *bbx14-3* (Wang *et al*., 2015). The specific guide RNA (gRNA) was designed using the web tool CHOPCHOP (http://chopchop.cbu.uib.no/) and cloned into pHEE401-E as described (Wang *et al*., 2015; Wang *et al*., 2022), and Col-0 plants were transformed with the construct via floral dipping using *Agrobacterium tumefaciens* GV3101 (Clough & Bent, 1998). Positive transformants were selected in the first generation (T1) on plates containing 0.5X MS medium supplemented with 50 μg mL^−1^ hygromycin and 1% (w/v) sucrose. To select for homozygous *bbx14* mutants, the *BBX14* gene was sequenced in the surviving plants using primers listed in Supporting Information Table S1.

For overexpression of BBX14 in Col-0, the *AT1G68520* coding region was amplified from cDNA by PCR (see Supporting Information Table S1 for primer information). The PCR product was then cloned with GATEWAY technology into both pB7FWG2 and pAUL1, thus generating fusions with the enhanced GFP-tag (eGFP) and HA-tag, respectively, which were expressed under the control of the Cauliflower Mosaic Virus *35S* promoter. Both constructs were introduced into Col-0 plants by floral dipping (Clough & Bent, 1998).

### Generation of tagged GLK1 lines

To generate *Arabidopsis thaliana* Col-0 plants expressing GLK1-GFP from its endogenous promoter, the genomic locus of *GLK1* (including the 3867-bp promoter but excluding the stop codon) was amplified by PCR and subcloned into the pCR8/GW/TOPO TA cloning vector (Invitrogen). The insert was transferred into the binary destination vector pMDC107 by LR reaction (pGLK1:gGLK1-GFP::pMDC107). Col-0 plants were transformed as described above.

### Protein isolation and immunoblot analyses

Proteins were homogenized in 2× Laemmli sample buffer (120 mM Tris-HCl, pH 6.8, 4% SDS, 20% glycerol, 2.5% β-mercaptoethanol, 0.01% bromophenol blue) and processed as described in Supporting Information Methods S2.

### Chlorophyll fluorescence analysis

*In vivo* chlorophyll *a* fluorescence of whole plants was recorded using an ImagingPAM chlorophyll fluorometer (Walz GmbH, Effeltrich, Germany) as described previously (Garcia-Molina *et al*., 2020).

### Measurement of chlorophyll content

For chlorophyll extraction, approximately 100 mg of leaf tissue from 3-week-old plants was ground in liquid nitrogen in the presence of 80% (v/v) acetone. After removal of cell debris by centrifugation, chlorophyll absorption was measured spectrophotometrically. Pigment concentrations were calculated following Lichtenthaler (1987) and normalized to fresh weight.

### Data analysis

Statistically significant differences in relative mRNA expression levels were tested by applying one-way ANOVA with Tukey’s post-hoc HSD test (https://astatsa.com; version August 2021). The significance of differences in root and hypocotyl lengths, Fv/Fm and Y(II) was tested by two-way ANOVA, followed by Tukey’s multiple comparison test (as indicated in the Figure legends) using GraphPad Prism version 9.4.1 for Windows (GraphPad Software, www.graphpad.com). Samples were always compared with the respective wild-type (Col-0) sample within the same condition.

### ChIP-seq sample preparation and data analysis

14-day-old seedlings (pGLK1:gGLK1-GFP in Col-0) were processed as described in Supporting Information Methods S3.

### RNA sequencing (RNA-Seq) and data analysis

Total RNA from plants was isolated using Trizol (Invitrogen, Carlsbad, USA) and purified using Direct-zol™ RNA MiniPrep Plus columns (Zymo Research, Irvine, USA) according to the manufacturer’s instructions, and RNA integrity and quality were assessed by an Agilent 2100 Bioanalyzer (Agilent, Santa Clara, USA). Generation of RNA-Seq libraries and 150-bp paired-end sequencing was carried out on an Illumina HiSeq 2500 (Illumina, San Diego, USA) or Novaseq6000 system by Novogene Biotech (Beijing, China) or Biomarker Technologies GmbH (Münster, Germany), respectively, using standard Illumina protocols. Three independent biological replicates were used per genotype. RNA-Seq reads were analyzed on the Galaxy platform (Afgan *et al*., 2016) essentially as described before (Xu *et al*., 2019), except that reads were mapped with the gapped-read mapper RNA STAR (Dobin *et al*., 2013).

## RESULTS

### Overexpression of BBX14 is deleterious

To gain insight into the function of BBX14, the mutant *bbx14-1* (SAIL_1221_D02) was identified in the SIGnAL database (Alonso *et al*., 2003); Fig. **S1a, b**), and found to express*BBX14* transcripts in amounts equivalent to 10% of Col-0 levels (Fig. **S1c**). Because *bbx14-1* was the sole T-DNA mutant line available, two independent CRISPR/Cas-mediated lines (*bbx14-2* and *bbx14-3*) were also generated. In *bbx14-2*, an insertion after nucleotide (nt) 924 relative to the start codon introduced a premature stop, and in *bbx14-3* a T-to-G change at nt 925 resulted in the replacement of a Trp by a Gly residue. In these lines, *BBX14* transcript levels were reduced to 10 and 25%, respectively, of that in Col-0 (Fig. **S1c**). To create plants that overexpress BBX14 (oeBBX14), Col-0 plants were transformed with a DNA fragment comprised of the *35S* Cauliflower Mosaic Virus (CMV) promoter and the coding sequence of *BBX14*, fused upstream of either the enhanced green fluorescence protein (eGFP) or a hemagglutinin (HA) tag. After selection for the resistance marker, 10 lines for each construct were identified, but fusion proteins were detected in only three of the oeBBX14-eGPF lines and two oeBBX14-HA lines (Fig. **S1d**). Only one of these lines expressed higher levels (2.2-fold) of the *BBX14* transcript than Col-0 plants, while the others contained slightly less than the wild-type control (Fig. **S1e**), suggesting that overexpression of BBX14 could be deleterious. To test this, the remaining seeds of the Col-0 plants bearing the overexpression constructs were again subjected to selection, and all resistant seedlings were transferred to MS. In this way, we identified lines that were paler and smaller than Col-0 seedlings (Fig. **1a**). Western blot analysis indicated that the severity of the seedling phenotype might be related to the degree of BBX14 overexpression (Fig. **1b**).

**Fig. 1.**
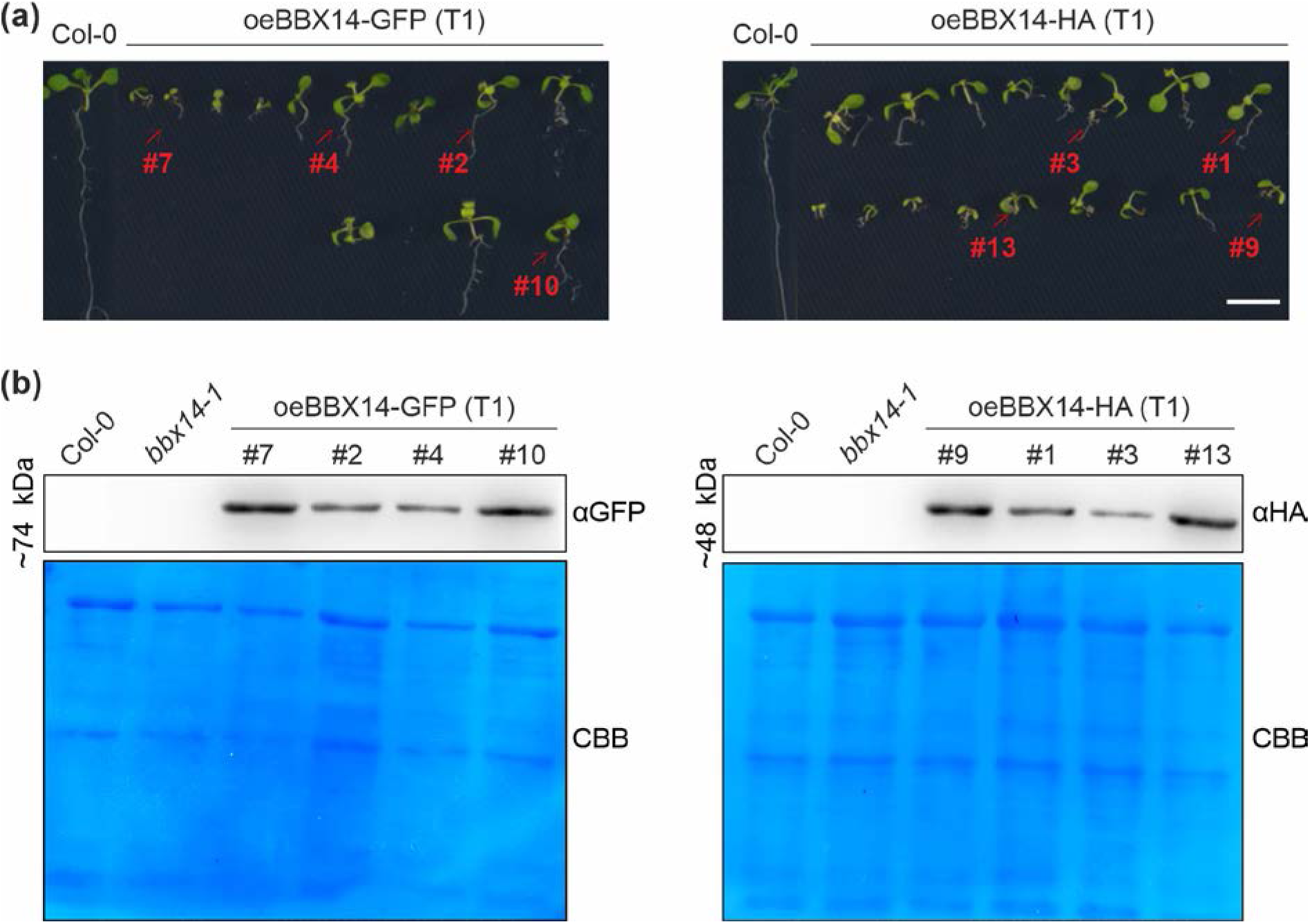
Overexpression of *BBX14* is deleterious. **(a)** Phenotypes of 10-day-old wild-type (Col-0) seedlings, and Col-0 seedlings transformed with constructs containing the coding region of *BBX14* fused to either the GFP- (left panel) or the HA-tag (right panel), which were placed under the control of the *35S* promoter. Red arrows indicate seedlings that provided the protein extracts that were subjected to SDS-PAGE. Scale bar = 0.5 cm. **(b)** Aliquots of total leaf proteins were isolated from plants as indicated in **(a)**, fractionated on SDS-PA gels (10%) under reducing conditions, and subjected to immunoblotting using antibodies raised against the GFP- or HA-tag, respectively. PVDF membranes were stained with Coomassie brilliant blue (CBB) to show protein loading.

### BBX14 is needed to acclimate plants to high light stress

Steady-state amounts of the *BBX14* transcript are reduced in plants exposed to high light (HL) levels (Kleine *et al*., 2007; Leister & Kleine, 2016; Huang *et al*., 2019; Garcia-Molina *et al*., 2020), as confirmed by the Genevestigator perturbations tool (https://genevestigator.com; Fig. **2a**). Interestingly, among all the *BBX* members represented on the Affymetrix ATH1 chip, this decrease in expression levels after HL treatment was quite specific for *BBX14*. In fact, Huang et al. (2019) identified *BBX14* as one of the top three hub genes in an HL co-expression network, suggesting that BBX14 might be an important regulator of the HL response. To test this, 1-week-old Col-0, *bbx14-1*, *bbx14-2* plants grown under normal lighting conditions (80 µmol photons m^-2^s^-1^) were exposed to a higher light level (1000 µmol photons m^-2^s^-1^), which was sufficient to strongly reduce BBX14 mRNA levels (Fig. **2b**). Importantly, for this experiment LED chambers were used which allowed for strict temperature control and heat contribution could be excluded. Maximum quantum yield of PSII (F_v_/F_m_) was monitored under control conditions, after 3, 8 and 12 h of HL treatment, and thereafter recorded after 12 and 36 h of de-acclimation. Under standard light levels, 1-week-old *bbx14* mutant plants largely resembled the WT (Fig. **2c**). Under HL treatment, a reduction of Fv/Fm (relative to WT) was readily apparent after 3 h (Fig. **2d**), and was intensified throughout the HL time course. However, *bbx14* plants could recover their Fv/Fm values after de-acclimation in normal growth conditions (Fig. **2c,d**).

**Fig. 2.**
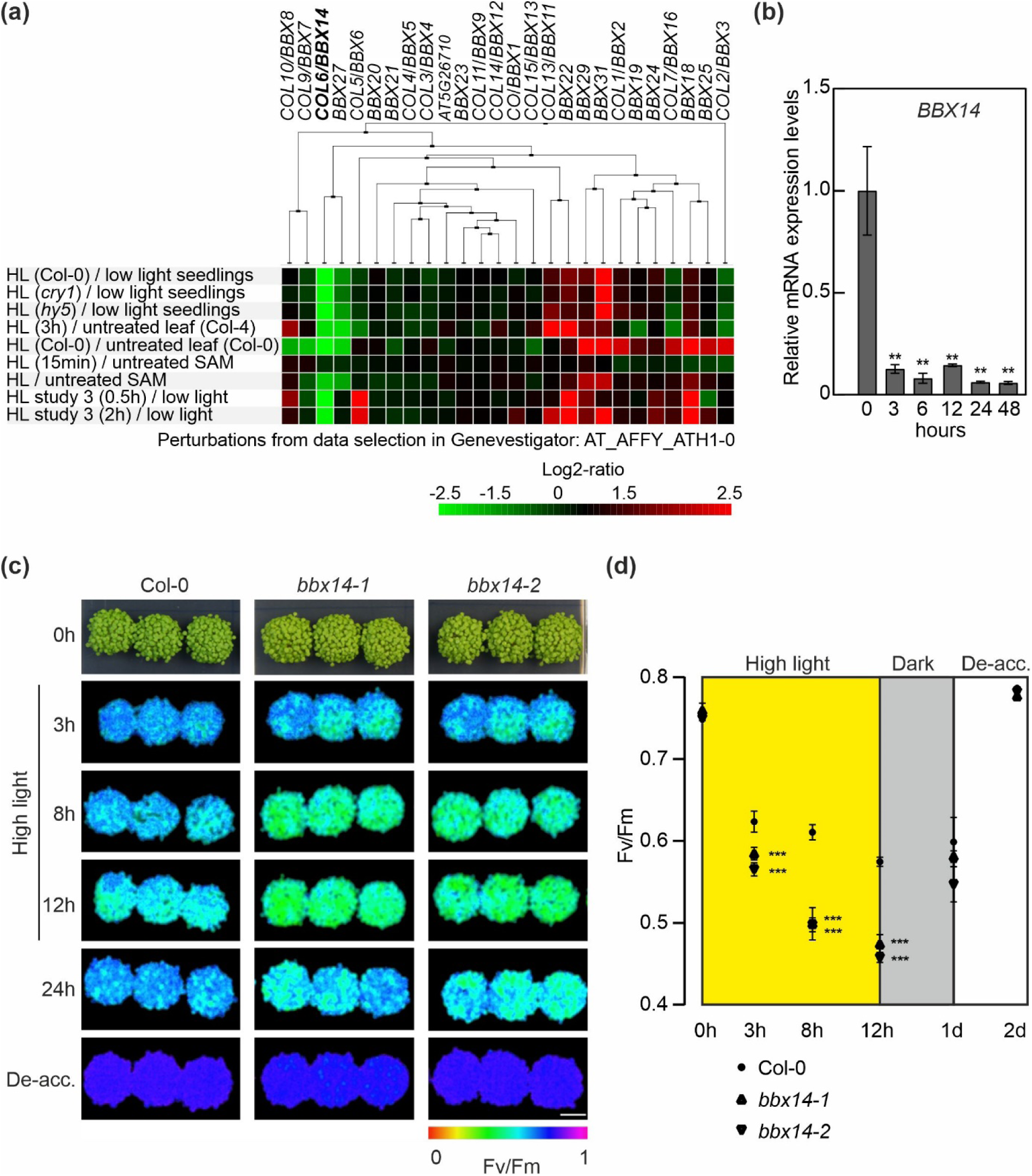
BBX14 is involved in high light (HL) acclimation. **(a**) Global profiling of *BBX* mRNA levels in response to perturbations was carried out with Genevestigator, and studies involving HL treatments are shown. The cladogram at the top summarizes the degree of relatedness between the expression profiles of the different *BBX* genes. SAM, shoot apical meristem. **(b)** RT-qPCR of *BBX14* expression in 7-day-old Col-0 plants grown under control conditions and then shifted to light level for up to two days. The results were normalized to *AT4G36800*, which encodes a RUB1-conjugating enzyme (RCE1). Expression values are reported relative to the corresponding transcript levels in Col-0, which were set to 1. Mean values ± SE were derived from two independent experiments, each performed with three technical replicates per sample. Statistically significant differences (Tukey’s test; \*\**P* < 0.01) between control and each HL time point are indicated. **(c)** Phenotypes and Imaging PAM pictures of Col-0 and mutant (*bbx14-1, bbx14-2*) plants grown for 1 week under control conditions (16-h light/8-h dark, 80 µmol photons m^−2^ s^−1^; left panel), shifted to high light (HL) conditions (16-h light/8-h dark, 1000 µmol photons m^−2^ s^−1^), and then de-acclimated (de-acc.) in control conditions. Scale bar = 1 cm. **(d)** Photosystem II maximum quantum yield (Fv/Fm) of wild-type (Col-0) and mutant (*bbx14-1 and bbx14-2*) seedlings grown as described in **(c)**.

Hence, BBX14 is beneficial for plant growth under HL stress.

### BBX14 is involved in seedling photomorphogenesis

BBX proteins have already been recognized as prominent factors in seedling photomorphogenesis (see Introduction). The involvement of BBX14 in this process is attested by the significantly longer hypocotyls seen in *bbx14* mutants compared to Col-0 when seedlings were grown for 3 days under long-day (LD) conditions (Fig. **3a**). Interestingly, the roots of *bbx14* mutants were also significantly longer than those of WT; in fact, this difference was even more pronounced than in hypocotyls. When seedlings were grown for 3 days in the dark, the *pifq* mutant, which lacks the phytochrome-interacting factors PIF1, −3, −4 and −5 and displays a partially constitutive photomorphogenic phenotype in the dark (Leivar *et al*., 2008), exhibited the expected phenotype, with a shorter hypocotyl and partially opened cotyledons (Fig. **3b**). The *bbx14* seedlings grown in the dark also had longer roots and hypocotyls than the WT, although this was true only for two of the three mutant lines examined (Fig. **3b**).

**Fig. 3.**
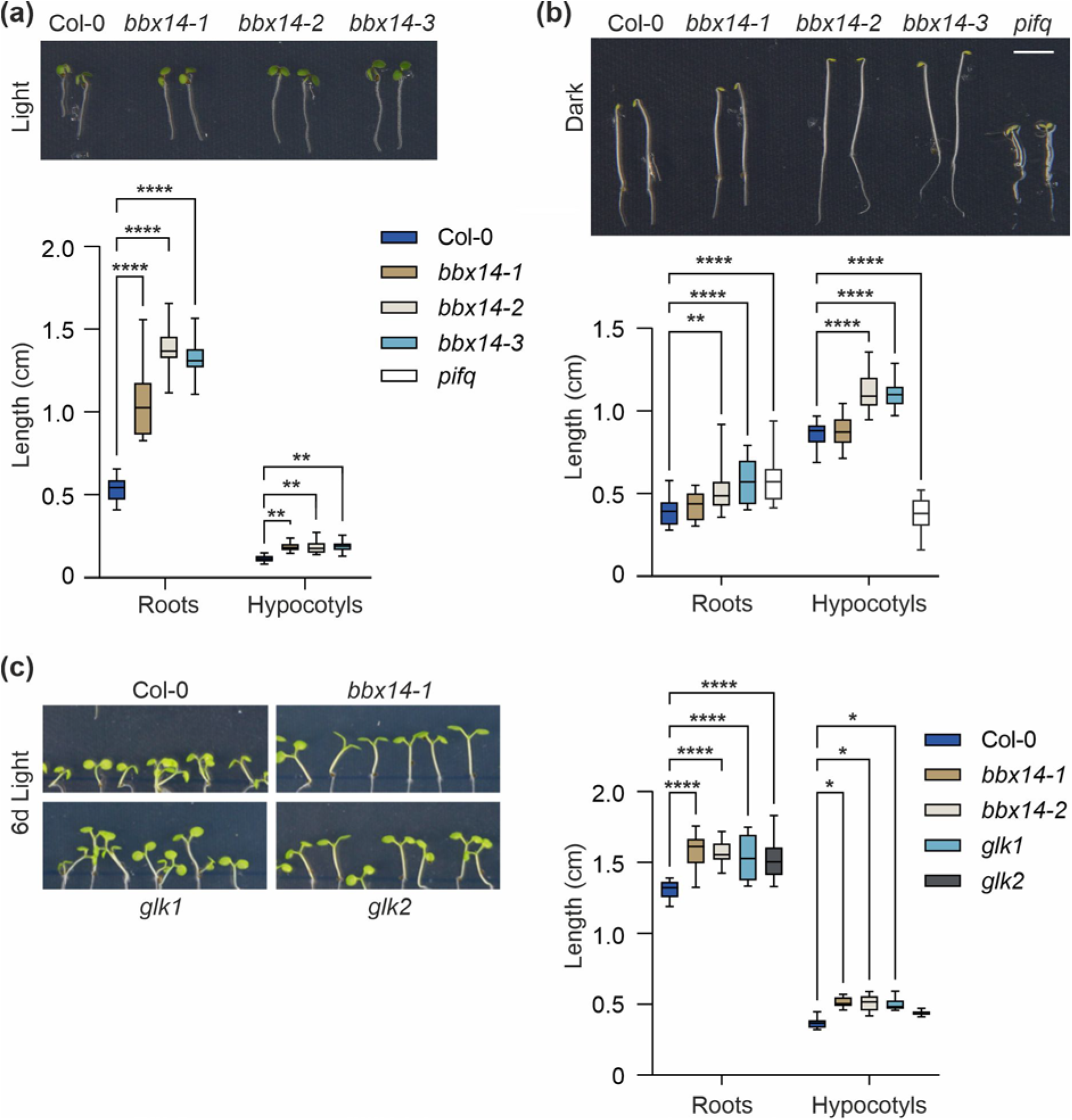
BBX14 is involved in photomorphogenesis. **(a)** Phenotypes (top) and quantification of root and hypocotyl lengths (bottom) of Col-0 and mutant (*bbx14-1, bbx14-2, and bbx14-3*) seedlings grown for 3 days under standard growth conditions (16-h light/8-h dark; 100 µmol photons m^−2^ s^−1^). **(b)** Phenotypes (top) and quantification of root and hypocotyl lengths (bottom) of Col-0 and mutant (*bbx14-1, bbx14-2, bbx14-3 and pifq*) seedlings grown for 3 days in the dark. **(c)** Phenotypes (top) and quantification of root and hypocotyl lengths (bottom) of Col-0 and mutant (*bbx14-1, bbx14-2 and glk1 and glk2*) seedlings grown for 6 days under standard growth conditions. (c, d, e) The center line of boxplots indicates the median, the box defines the interquartile range, and the whiskers indicate minimum and maximum values. Statistically significant differences between wild-type and each mutant line are indicated (two-way ANOVA; \*\*\*\**P* < 0.0001, \*\*\**P* < 0.001, \*\**P* < 0.01, \**P* < 0.05). Scale bar = 0.5 cm.

It has already been noted that levels of *BBX14* mRNA are dependent on GLK1 (Veciana *et al*., 2022), and *glk1* seedlings develop longer hypocotyls in the light. Accordingly, we found that hypocotyl lengths of *bbx14*, *glk1* and *glk2* mutants were likewise elongated (Fig. **3c**), and our data indicate that *glk1* seedlings also have a long-root phenotype similar to that caused by a lack of BBX14 under our growth conditions (Fig. **3c**). To the best of our knowledge, this has not been reported previously, but an involvement of GLKs in root growth is independently supported by the reduction in root lengths seen in oeGLK lines grown under phosphate limitation (Kang *et al*., 2014).

### The *BBX14* gene is a direct target of GLK1

Levels of *BBX14*, *BBX15* and *BBX27* transcripts are elevated in oeGLK1 and oeGLK2 lines (Waters et al., 2009), which suggests that these genes are targets of GLKs. Moreover, in light of the similarity between the *bbx14* and *glk1* hypocotyl-length phenotypes, we wanted to clarify whether or not *BBX14* is a direct target of GLK1. To this end, we performed a chromatin-immunoprecipitation experiment, followed by sequencing (ChIP-seq), on 14-day-old seedlings of a plant line that expresses GLK1 from its endogenous promoter in the Col-0 background. Our ChIP-seq experiment confirmed binding of GLK1 to the *BBX16* promoter (Veciana *et al*., 2022), and detected a GLK1-bound genomic region directly upstream of the *BBX14* transcription start site (TSS). Analysis of this region revealed the presence of four GLK1 binding sites (Fig. **4a**), all of which match the CCAATC consensus identified in a set of co-expressed, photosynthesis-related genes (Kobayashi *et al*., 2012). One of these motifs, located 70 bp upstream of the TSS, could be further extended to yield a sequence that matches a GLK1-binding motif identified by protein-binding microarrays (**GATTCT**GATTGG; (Franco-Zorrilla *et al*., 2014) and thus represents a strong candidate for GLK1 binding. This result indicates that *BBX14* is indeed a direct target of GLK1, and additional *BBX* genes were detected among the top 10 most highly enriched potential target genes (Supporting Information Table S2; Fig. **4b**). Furthermore, *TIME FOR COFFEE* (*TIC*), a component of the circadian clock, was among the top targets. Besides a motif that was identified as “G2-like” but resembles a GLK1 binding consensus, MEME identified motifs resembling bZIP, LOB, C2H2 and GATA TF binding motifs as markedly enriched in the GLK1-bound genomic regions (Fig. **4c**), a finding that implies coordinated control of target gene activities. In fact, the *BBX14* core promoter is bound by GLK1 in combination with other TFs involved in light/circadian and cytokinin signaling, such as PIF4, LUX, HY5, PRR5 and ARR10 based on the ChIP-hub platform (Fu *et al*., 2022).

**Fig. 4.**
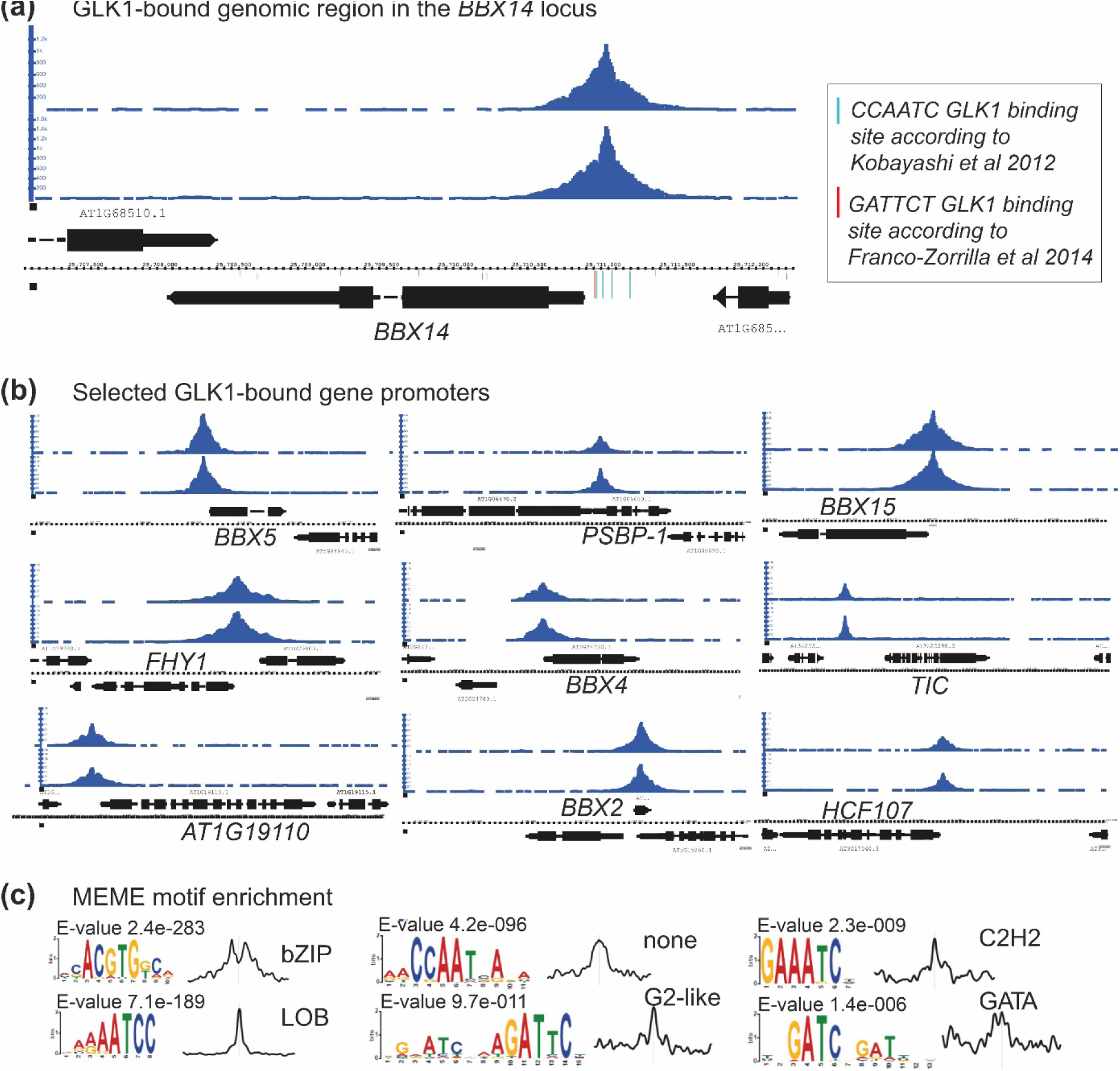
*BBX14* is a target of GLK1. **(a)** Chromatin-immunoprecipitation followed by sequencing (ChIP-seq) was performed with 14-day-old seedlings of a plant line that expresses GLK1 from its endogenous promoter in the Col-0 background (*pGLK1*:*GLK1*-*GFP*), and a snapshot of the *BBX14* gene region is shown. **(b)** Snapshots of selected genes targeted by GLK1. **(c)** Additional motifs bound by GLK1 identified by the MEME Suite (Bailey *et al*., 2015).

### Transcriptomes of etiolated *bbx14* and *glk1 glk2* seedlings

To identify putative target genes of BBX14, we first looked for transcriptome changes, focusing on light-dependent and -independent or common targets. RNAs were isolated from 3-day-old etiolated WT and *bbx14-1* seedlings, and from 3-day-old etiolated seedlings that had been shifted to LD conditions for 1 day, and subjected to RNA sequencing (RNA-Seq). GLKs are also required for chloroplast development in the absence of light, as shown by lower accumulation of the chlorophyll precursor protochlorophyllide (Pchlide) in the *glk1 glk2* mutant (Waters *et al*., 2009). Since to our knowledge no transcriptomic data for dark-grown *glk1 glk2* seedlings have been generated up to now, RNAs isolated from 3-day-old etiolated *glk1 glk2* seedlings were included.

In dark-grown *bbx14-1* seedlings, only four transcripts were reduced and two elevated relative to WT (>2-fold, *P* < 0.5; Supporting Information Table S3). Levels of three of the former (*ARP9*, *ARABIDOPSIS RESPONSE REGULATOR 7* (*ARR7*) and *AT5G26270*), were also reduced in dark-grown *glk1 glk2* seedlings (Fig. **5a,b**). In the *glk1 glk2* mutant, expression levels of 83 and 184 genes were significantly reduced and elevated, respectively (Fig. **5a,b**; Supporting Information Table S4). Among the up-regulated set, gene ontology (GO) analysis with DAVID (Sherman *et al*., 2022) identified “biological process (BP)” and “cellular response to sucrose starvation” as the most highly enriched GO categories (Fig. **5c**). One representative of the latter category is *AT3G47340*, which codes for GLUTAMINE-DEPENDENT ASPARAGINE SYNTHASE 1 (ASN1)/DARK INDUCIBLE6 (DIN6), and was also identified as a direct target of GLK1 (Supporting Information Table S2). Expression of *DIN6* (a target for the Snf1-related kinase SnRK1) is induced by conditions that limit photosynthesis and respiration, and hence affect sugar and energy supply (Baena-Gonzalez *et al*., 2007). This may imply that sugar supplies are lower in dark-grown *glk1 glk2* seedlings. Among the down-regulated genes in *glk1 glk2*, DAVID detected remarkably highly enriched (251-, 254- and 109-fold, respectively) GOs in the categories “cellular component” (CC), BP and “molecular function” MF, all of which are associated with the light-harvesting complex of PSII (Fig. **5c**). The drastic reduction (to 4.5%) in levels of expression of the retrograde signaling marker gene *LHCB1.2*, which codes for light-harvesting chlorophyll a/b-binding proteins of photosystem (PS) II, is particularly striking, and is even more marked than that seen (8.3%) when *glk1 glk2* plants were grown under LD conditions (Bieluszewski *et al*., 2022). Moreover, expression of *BBX14* is not only dependent on GLKs in the light, since *BBX14* RNA levels are readily down-regulated to half of WT levels in dark-grown *glk1 glk2* seedlings.

**Fig. 5.**
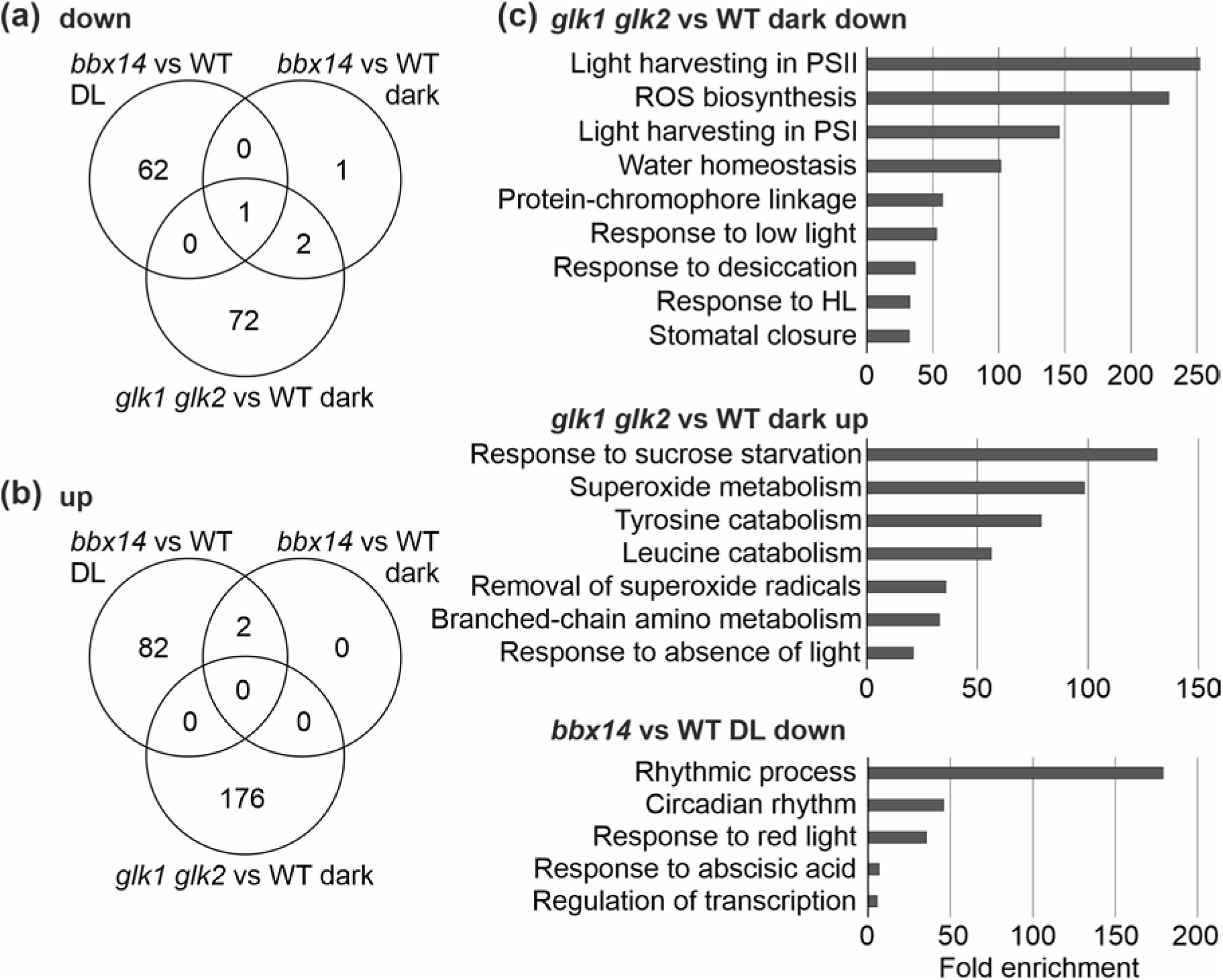
Comparative analysis of transcriptome changes in *bbx14* and *glk1 glk2* mutant seedlings. Venn diagrams depicting the degree of overlap between the sets of genes whose expression levels were reduced **(a)** or elevated **(b)** by at least two-fold in *bbx14-1* seedlings that had been exposed to LD conditions and dark-grown *bbx14-1* and *glk1 glk2* seedlings compared with their respective WT (Col-0) control. **(c)** GO analysis of genes whose expression is changed in light-shifted *bbx14-1* seedlings and dark-grown *glk1 glk2* seedlings compared to their respective WT control. GO annotations for the biological process category were extracted from DAVID. For *glk1 glk2* versus WT only more than 20-fold enriched terms are shown.

In addition, 30 of the 109 genes that were down-regulated in dark-grown *glk1 glk2* seedlings were identified as targets of GLK1. In contrast, only 23 of the 332 genes that were up-regulated were bound by GLK1 (Supporting Information Tables S2 and S3), which supports the idea that GLK1 mainly acts as a transcriptional activator.

### BBX14 participates in the regulation of genes associated with the circadian clock

In the *bbx14-1* seedlings that had been exposed to LD conditions for a day, 147 transcripts changed significantly relative to WT (>2-fold, *P* < 0.5; Supporting Information Table S5), of which 63 were reduced and 84 were elevated (Fig. **5a,b**). Of these, one (*ACTIN-RELATED PROTEIN 9* (*ATARP9*) and two, respectively, were among those whose expression was altered in the same sense in etiolated *bbx14* seedlings, and therefore represent prime targets of BBX14. Moreover, transcript levels of the clade II members *BBX7* and *BBX8* were reduced. Lower levels of *PHOTOPERIODIC CONTROL OF HYPOCOTYL 1* (*PCH1*) and higher levels of *XYLOGLUCAN ENDOTRANSGLUCOSYLASE/HYDROLASE 9* (*XTH9*) and *XTH16* transcripts might account for the longer hypocotyls, because XTHs encode proteins involved in the weakening of cell-cell contacts and rearrangement of the cell wall. The GO “rhythmic process” was detected with a striking 180-fold enrichment in the set of genes with reduced transcript levels (Fig. **5c**), represented for example by genes encoding the flavin-binding kelch-repeat F-box protein FKF1, PSEUDO-RESPONSE REGULATOR 5 (PRR5) and the CCT motif-containing response regulator protein TIMING OF CAB EXPRESSION 1 (TOC1). These findings suggest a role for BBX14 in the circadian clock, as has already been shown for BBX19 (Yuan *et al*., 2021). The binding of circadian clock-associated TFs to the *BBX14* promoter (see above) suggests the presence of feedback regulatory mechanisms, but here it has to be noted that, although *BBX14* expression is diurnally regulated, it was not detected as a strongly rhythmically active gene by the Arabidopsis eFP Browser (Winter *et al*., 2007).

### The *bbx14* hypocotyl phenotype depends on a retrograde signal

In germinating seedlings, chloroplast biogenesis is highly sensitive to environmental changes and requires ‘biogenic control’ (Pogson *et al*., 2008), i.e. signaling from chloroplasts to the nucleus during early developmental stages in which a GLK1-GUN1 module is involved (Leister & Kleine, 2016; Martin *et al*., 2016). When WT seedlings are grown under very low levels of white light (1 µmol m^-2^ s^-1^), LIN partially prevents de-etiolation, and hypocotyls are longer than those of seedlings grown in its absence (Martin *et al*., 2016). Because *bbx14* seedlings displayed a longer hypocotyl phenotype under our light intensity (100 µmol m^-2^ s^-1^) – a condition which serves as a test for *gun* phenotypes – we assessed hypocotyl growth in WT, *bbx14* and *gun1-102* lines in the presence of LIN. Under our lighting conditions, hypocotyl lengths of WT seedlings were independent of LIN supplementation (Fig. **6a**), indicating that the etiolation phenotype in the presence of LIN is mainly observed under very low light levels. This is compatible with the fact that, when 2-day-old etiolated seedlings were transferred to white-light levels of 25 µmol m^-2^ s^-1^, the same hypocotyl lengths were observed in the absence and presence of LIN (Martin *et al*., 2016). Strikingly, the long hypocotyl (and also root) phenotype of *bbx14*, *glk1* and *glk2* seedlings was alleviated when seedlings were grown on LIN, and hypocotyl lengths were comparable to those of the WT (Fig. **6b**), which suggests that the *bbx14*, *glk1* and *glk2* hypocotyl phenotypes are dependent on a retrograde signal. It was previously shown that the retrograde signaling and phytochrome pathways antagonistically regulate the PIF-repressed transcriptional network (Martin *et al*., 2016), and the core promoter of *BBX14* is bound by PIF4 in addition to GLK1 (see above). Notably, *BBX14* mRNA levels were de-repressed in dark-grown *pifq* seedlings, and lincomycin treatment prevented this de-repression (Fig. **6c**).

**Fig. 6.**
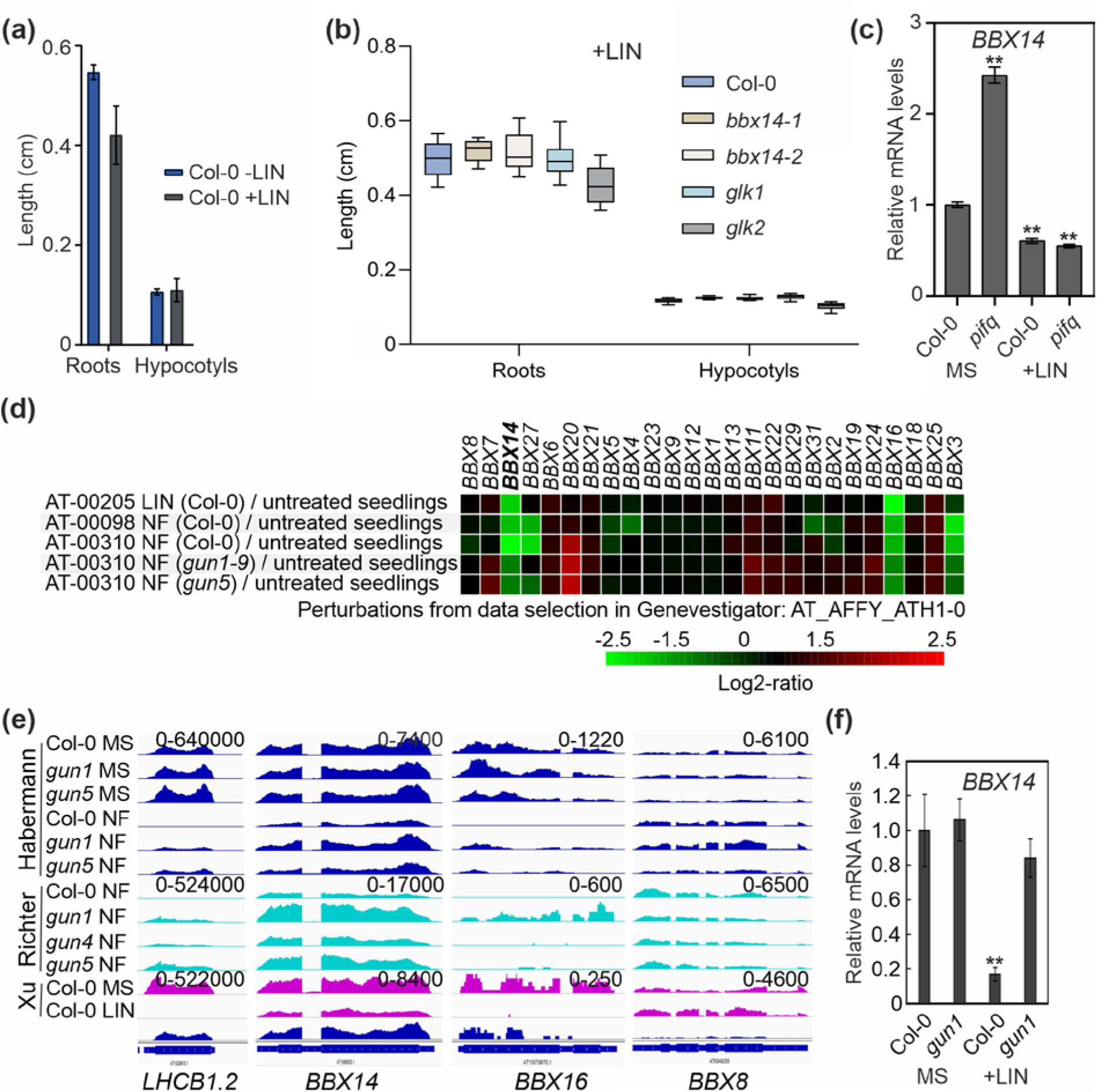
The *bbx14* hypocotyl phenotype and *BBX14* mRNA expression depend on retrograde signaling. **(a)** Quantification of root and hypocotyl lengths of Col-0 seedlings grown undert standard growth conditions (16-h light/8-h dark and 100 µmol photons m^−2^ s^−1^) in the absence (-LIN) or presence (+LIN) of lincomycin. Bars represent averages, while error bars represent standard errors. **(b)** Quantification of root and hypocotyl lengths of 6-day-old wild-type (Col-0) and mutant (*bbx14-1, bbx14-2, glk1 and glk2)* seedlings grown under standard growth conditions in the presence of lincomycin (+Lin). **(c)** RT-qPCR of *BBX14* mRNA levels in 3-day-old dark-grown Col-0 and *pifq* seedlings in the absence (MS) or presence of lincomycin (+LIN). RT-qPCR was performed as described in the legend to Fig. 2b. Expression values are reported relative to the corresponding transcript levels in Col-0 grown on MS, which were set to 1. **(d)** The reduction of *BBX14* mRNA levels during biogenic signaling depends on GUN1. Global profiling of *BBX* mRNA levels in response to perturbations were determined with Genevestigator, and studies involving lincomycin (LIN) and norflurazon (NF) treatments are shown. **(e)** Snapshots of re-analyzed RNA-Seq data published by (Habermann *et al*., 2020), (Richter *et al*., 2020) and (Xu *et al*., 2020). The read depths were visualized with the Integrative Genomics Viewer (IGV). **(f)** RT-qPCR of *BBX14* mRNA levels in Col-0 and *gun1* seedlings grown for 4 days in continuous light (100 µmol photons m^−2^ s^−1^) in the absence (MS) or presence of lincomycin (+LIN). **(f)** Expression values are reported relative to the corresponding transcript levels in Col-0 grown on MS, which were set to 1. Mean values ± SE were derived from three independent experiments, each with three technical replicates. Statistically significant differences (Tukey’s test; \*\**P* < 0.01) between wild-type and mutant are indicated.

### Reduction of *BBX14* mRNA levels during biogenic signaling is dependent on GUN1

Data extracted from Genevestigator (https://genevestigator.com) suggested that, of the *BBX* members represented on the Affymetrix ATH1 chip, levels of *BBX3*, *BBX14*, *BBX16* and *BBX27* mRNAs are reduced in NF-treated seedlings, while under LIN conditions only *BBX14* and *BBX16* mRNAs are partially repressed (Fig. **6d**). As a complement to Genevestigator results, previously published RNA-Seq data (Habermann *et al*., 2020; Richter *et al*., 2020; Xu *et al*., 2020) were re-analyzed and the read depths of *BBX14* and *BBX16* transcripts were visualized with the Integrative Genomics Viewer (IGV; Fig. **6e**). As controls, read depths of *LHCB1.2* and *BBX8* were plotted. *LHCB1.2* mRNA levels are known to be strongly reduced under NF or LIN treatment (Oelmuller *et al*., 1986), and *BBX8* was used as an example of other BBX members whose RNA levels were suggested not to be reduced under LIN or NF treatment. All investigated loci behaved as expected, and confirmed the Genevestigator data (Fig. **6e**). Moreover, these data suggested that the decrease in *BBX14* and *BBX16* mRNA levels under NF conditions is alleviated in *gun1* and *gun5* mutants. BBX16 has previously been implicated in retrograde signaling (Veciana *et al*., 2022), and we suspected that BBX14 might also be a component of the GUN1-GLK1 module. The *gun1* mutant is special in that – unlike all the other *gun* mutants – it also shows the *gun* phenotype under LIN conditions (summarized in (Richter *et al*., 2023). To find out whether LIN-mediated repression of *BBX14* is also dependent on GUN1, *BBX14* mRNA expression was evaluated in 4-day-old Col-0 and *gun1-102* seedlings grown on MS in the absence or presence of LIN. Under LIN treatment *BBX14* levels were reduced to 17% in Col-0 compared to control conditions (Fig. **6f**), and this effect was dependent on GUN1 since *BBX14* expression recovered almost completely in *gun1-102* seedlings treated with LIN. This, together with ChIP-Seq data (see above) and the nuclear localization of BBX14 (Fig. **S2**), suggests that BBX14 might act as a receptor for GUN1/GLK1-dependent retrograde signals in the nucleus.

### Involvement of BBX14 in GUN-mediated retrograde signaling

We hypothesized that, like oeGLK lines (Leister & Kleine, 2016; Martin *et al*., 2016), overexpression of BBX14 could result in a *gun* phenotype, or a lack of BBX14 could evoke a *LHCB1.2* hypersensitive phenotype in the presence of LIN. To test this, Col-0, *gun1-102* (as a control), the *bbx14* mutant and the strongest “overexpression” seedlings were grown in the presence or absence of LIN for 4 days under continuous light (100 µmol m^−2^ s^−1^). After LIN treatment, *gun1-102* showed, as expected, enhanced expression of *LHCB1.2* mRNA relative to WT seedlings, but *LHCB1.2* levels were unchanged in both *bbx14* mutants and the “oeBBX14” line (Fig. **S3a**). Also, the expression of *LHCB2.1*, *LHCB2.4* was not affected in lines with altered *BBX14* levels (Fig. **S3b**). However, slightly higher *CA1* levels were observed in “oeBBX14” (Fig. **S3b**). It is reasonable to suppose that 2.2-fold induction of BBX14 (see Fig. S1) is insufficient to cause a true *gun* phenotype, so we made use of an inducible BBX14 line (named TPT14; only one line available) generated within the TRANSPLANTA collection (Coego *et al*., 2014). After 4 days of induction, TPT14 seedlings were clearly perturbed, while they looked healthy when not induced (Fig. **S4a**), in agreement with the effects of stable overexpression of BBX14 (see Fig. 1). Notably, overexpression of BBX14 resulted in the shortening of roots and hypocotyls (Fig. **S4b,c**). Hence, in order to circumvent secondary effects, we employed a different experimental set-up to test for *gun* phenotypes. seedlings were first grown for three days in darkness in the absence or presence of inhibitor (NF or LIN), then sprayed with the inducer, put back in the dark for 2 h, transferred into the light for 16 h (100 µmol m^-2^ s^-1^), and then harvested for RT-qPCR. Interestingly, *BBX14* induction was very successful when seedlings were treated with NF, but in seedlings treated with LIN or grown without inhibitor, induction was weaker (Fig. **7a**, S**5a**). In the absence of inhibitor or on LIN, *LHCB1.2*, *CA1* and *RBCS1A* expression levels were similar to those seen in Col-0 in all lines tested, apart from slightly elevated *LHCB1.2* levels in *gun1* and *TPT14* seedlings grown without inhibitor (Fig. **S5b**). In addition, the suitability of this protocol for *gun* phenotype evaluation was confirmed by de-repression of the retrograde signaling markers *LHCB1.2*, *CARBONIC ANHYDRASE 1* (*CA1)* and *RIBULOSE BISPHOSPHATE CARBOXYLASE SMALL CHAIN 1A* (*RBCS1A*) in *gun1* compared to Col-0 when seedlings were grown on NF (Fig. **7b**). In response to NF, *LHCB1.2* mRNA levels were only slightly elevated in TPT14 compared to Col-0 (Fig. **7b**), but interestingly, those of *CA1* and *RBCS1A* were comparable to levels found in the *gun1* mutant in the TPT14 line (Fig. **7b**).

**Fig. 7.**
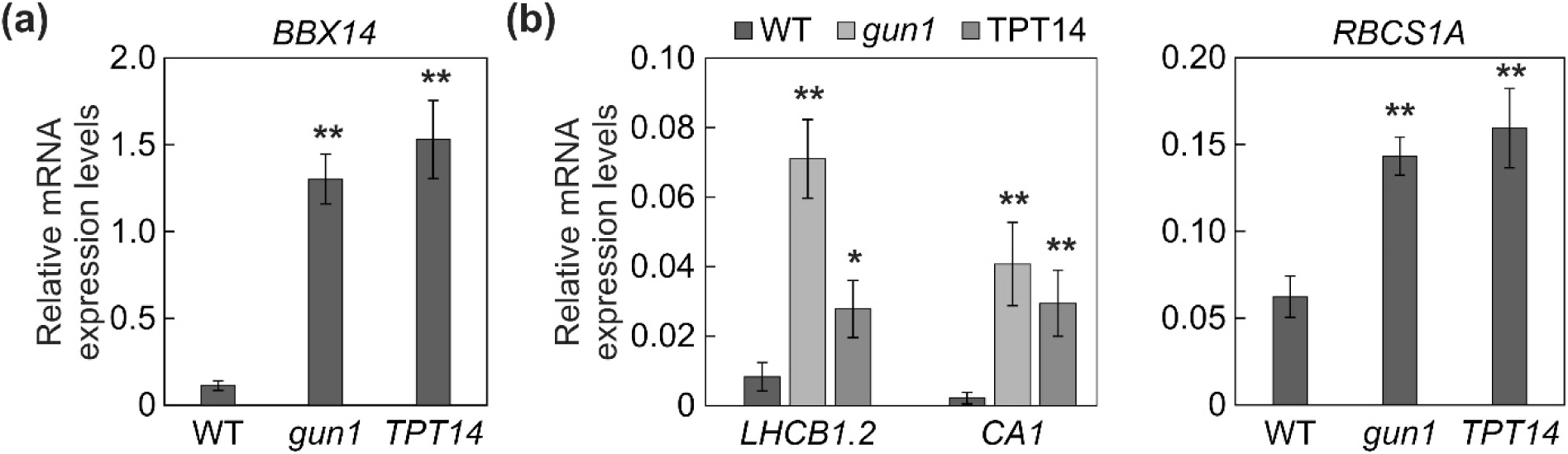
Overexpression of BBX14 confers a partial *gun* phenotype. **(a)** Levels of *BBX14* mRNAs in seedlings grown for three days in the dark in the presence of norflurazon (NF), sprayed with inducer, put back for 2 h in the dark, placed for 16 hours into light (100 µmol m^-2^ s^-1^) and then harvested for RT-qPCR. **(b)** Transcript levels of retrograde marker genes *LHCB1.2*, *RBCS1A* and *CA1* of seedlings grown on NF and treated as in **(a)**. **(a)**, **(b)** The results were normalized to *RCE1*. Expression values are reported relative to the corresponding transcript levels in Col-0, which were set to 1. Mean values were derived from four independent experiments, each with three technical replicates. Bars indicate standard deviations. Statistically significant differences (Tukey’s test; \*\**P* < 0.01; \**P* < 0.05) between Col-0 and mutant samples are indicated.

These results suggest that, like BBX16 (Veciana *et al*., 2022), BBX14 does not mediate regulation of *LCHB1.2* during retrograde signaling, but does alter the expression of other *PhANG*s, such as *CA1* and *RBCS1A*.

## DISCUSSION

BBX proteins are vital for plant development, and have also emerged as players in the process of acclimation to adverse environmental conditions (Alvarez-Fernandez *et al*., 2021; Talar & Kielbowicz-Matuk, 2021). Together with BBX15-BBX17, BBX14 belongs to clade III of the B-box proteins (Khanna *et al*., 2009). BBX16 was first described 10 years ago as a phytochrome-B-dependent regulator of branching and the shade avoidance response (Wang *et al*., 2013; Zhang *et al*., 2014). Meanwhile, more information has become available about the functions of the members of this clade. Thus, BBX17 was recently suggested to interact with CONSTANS to negatively regulate flowering time (Xu *et al*., 2022), BBX14/15/16 were identified as components of a GLK-BBX module that inhibits precocious flowering (Susila *et al*., 2023), and BBX16 has emerged as a promoter of seedling photomorphogenesis and as an element that acts downstream of the GUN1/GLK1 module during retrograde signaling (Veciana *et al*., 2022). Indeed, BBX14 first aroused our interest when we identified it in a core-response module for the high-light (HL) and retrograde signaling response (Leister & Kleine, 2016) which at the interface of retrograde and acclimation pathways.

### The role of BBX proteins in acclimation to light stress

Alvarez-Fernandez *et al*. (2021) defined an accelerated HL acclimation phenotype as a significant enhancement of the operating efficiency of photosystem II (PSII; Y(II)). Because *BBX14* is co-regulated with *PhANG* transcripts under HL conditions (Huang *et al*., 2019; Garcia-Molina *et al*., 2020), this could in principle be achieved if higher BBX14 levels increased transcription levels of *PhANG* genes under HL, or a lack of BBX14 would result in less high-light tolerant plants, which was indeed the case (see Fig. 2).

Along with *BBX14*, transcriptional levels of all other clade III members and those of *BBX27* have been found to decrease under HL, while other *BBX* transcripts, including those of the clade V members *BBX29*-*BBX32*, were elevated (Huang *et al*., 2019; Garcia-Molina *et al*., 2020). A *bbx32* mutant displayed slightly higher Y(II) after 5 days of HL treatment, while plants overexpressing *BBX32* were impaired in acclimation to HL. But overexpression of BBX32 was already slightly detrimental under growth conditions with reduced amounts of several *PhANG* transcripts (Alvarez-Fernandez *et al*., 2021). In contrast to BBX14, the functions of BBX32 have already been well described, and the characterization of this protein is a prime example of the complex regulatory and signal transduction cascades in which BBX proteins are involved, including redundancies, feedback regulations, opposing and multi-layered functions. Therefore, seedlings that overexpress *BBX32* develop elongated hypocotyls and are hyposensitive to various light conditions, whereas *bbx32* mutants exhibit short hypocotyls under low-fluence conditions. BBX32 acts as a repressor of photomorphogenesis by interacting with BBX21 and suppressing its transcriptional activity in HY5-dependent and -independent pathways (Holtan *et al*., 2011). Moreover, BBX32 interacts with BRASSINAZOLE-RESISTANT 1 (BZR1) and PIF3, and is suggested to integrate light and brassinosteroid signals to regulate cotyledon opening during de-etiolation (Ravindran *et al*., 2021), On the other hand, *BBX20* is transcriptionally repressed by BZR1 (Fan *et al*., 2012). Finally, in the presence of BBX32, BBX4 binds to the promoter of *FLOWERING LOCUS T* (*FT*) to repress *FT* expression and flowering. However, both overexpression and silencing (microRNA) of *BBX32* results in late flowering (Tripathi *et al*., 2017).

Overall, a picture is emerging in which BBX proteins have multi-layered functions in plant development, as well as playing roles in responses to acute stress.

### The function(s) of BBX clade III members in photomorphogenesis

As mentioned above, studies of BBX proteins are hampered by functional redundancies and/or feedback regulation. For example, the hypocotyls of the *bbx28 bbx29 bbx30 bbx31* quadruple mutant are shorter than those of double and single mutants (Song *et al*., 2020), which suggests that the clade V members BBX28, BBX29, BBX30 and BBX31 additively repress seedling photomorphogenesis (Song *et al*., 2020). Further complications arise from the fact that, while *BBX30* and *BBX31* are transcriptionally repressed by HY5 (Heng *et al*., 2019), BBX28 and BBX29 act together to prevent HY5 from binding to the promoters of *BBX30* and *BBX31,* which results in the expression of these genes. In addition, BBX30 and BBX31 interact with the promoters of *BBX28* and *BBX29* and enhance their expression (Song *et al*., 2020), and BBX29 undergoes COP1-mediated degradation in the dark (Heng *et al*., 2019). Within clade III, BBX15 is most closely related to BBX14, followed by BBX16 and BBX17 (Khanna *et al*., 2009). While BBX14’s function in seedling photomorphogenesis is clear (our results), its involvement in the regulation of flowering time was only revealed by the simultaneous knock-down of BBX14, - 15 and −16 (Susila *et al*., 2023). Conversely, cotyledon phenotypes were more clearly revealed by overexpression of BBX16 than in *bbx16* mutant lines (Veciana *et al*., 2022). BBX14-16 are all targets of GLK1, but it can be safely assumed that future research will uncover still more complex modes of action for these proteins. Such studies will also clarify whether BBX15 affects photomorphogenesis and whether a *bbx14 bbx15 bbx16* triple mutant has a greater impact on plant development than any of the single or double mutant combinations.

### BBX14 as a direct target of GLK1 in GUN1-dependent retrograde signaling

A screen designed to uncover components of the retrograde signaling pathway in seedlings led to the identification of the chloroplast-localized GENOMES UNCOUPLED (GUN) proteins (see Introduction (Susek *et al*., 1993). While all *gun* mutants de-repress *PhANG*s under NF treatment, *gun1* alone responds in the same manner in the presence of LIN (Koussevitzky *et al*., 2007). During GUN1-dependent signaling, the GLK1/2 proteins receive the signals in the nucleus (Leister & Kleine, 2016; Martin *et al*., 2016) which are key regulators of *PhANG* transcript accumulation (Waters *et al*., 2009). It has previously been noted that light-dependent and chloroplast signaling pathways converge at some point (Ruckle *et al*., 2007; Leister *et al*., 2014; Martin *et al*., 2016), and recently BBX16 was identified as such a component (Veciana *et al*., 2022). *BBX16* was shown to be a direct target of GLK1 by ChIP-qPCR (Veciana *et al*., 2022; Susila *et al*., 2023) and indeed, BBX16 mediates the expression of a subset of GLK1-regulated *PhANG* genes under LIN conditions (Veciana *et al*., 2022). To detect further targets of GLK1, we chose a genome-wide approach (ChIP-Seq) and identified *BBX14*, -*15* and *BBX2*, *-4* and *-5* as additional direct targets of GLK1 (see Fig. 4, and Supporting Information Table S2). They therefore fulfill the first criterion for membership of the GUN1/GLK-dependent biogenic signaling pathway. The second and third criteria are transcript reduction in the presence of both NF and LIN and functional dependence on GUN1, respectively. But of the BBX members that fulfill criteria 2 and 3 (see Fig. S6), only BBX14, - 15 and -16 also match criterion 1. Overexpression of BBX14 results in a *gun* phenotype similar to that seen in oeBBX16 lines. However, this result has to be viewed with caution, because we had only one inducible oeBBX14 line to hand and stable overexpression is too harmful for the plant. Branching of the signaling pathway downstream of GLK1, as discussed by (Veciana *et al*., 2022), might enable BBX14 to regulate a subset of target genes, and additive functions might play a role here. Whether clade only clade III BBX proteins have evolved to participate in retrograde signaling, or BBX factors from other clades are also involved remains to be determined.

In addition to branching of pathways, a complex interplay involving for example antagonistic regulatory mechanisms and feedback loops, such as those described above, should be considered. This holds for GLK1 regulation in response to plastid signals. Levels of the GLK1 transcript decline when plastid biogenesis is inhibited, and this response is GUN1-dependent (Kakizaki *et al*., 2009). In contrast, NF- or LIN-treated *gun1* mutant seedlings fail to accumulate the GLK1 protein, suggesting that plastid signals regulate GLK1 protein levels in a GUN1-independent manner. Reduced GLK1 protein levels in damaged plastids are partially restored by MG132, a proteasome inhibitor, indicating that the ubiquitin-proteasome system participates in the degradation of GLK1 in response to plastid signals (Tokumaru *et al*., 2017), which adds yet another layer of complexity (Fig. **8**).

**Fig. 8.**
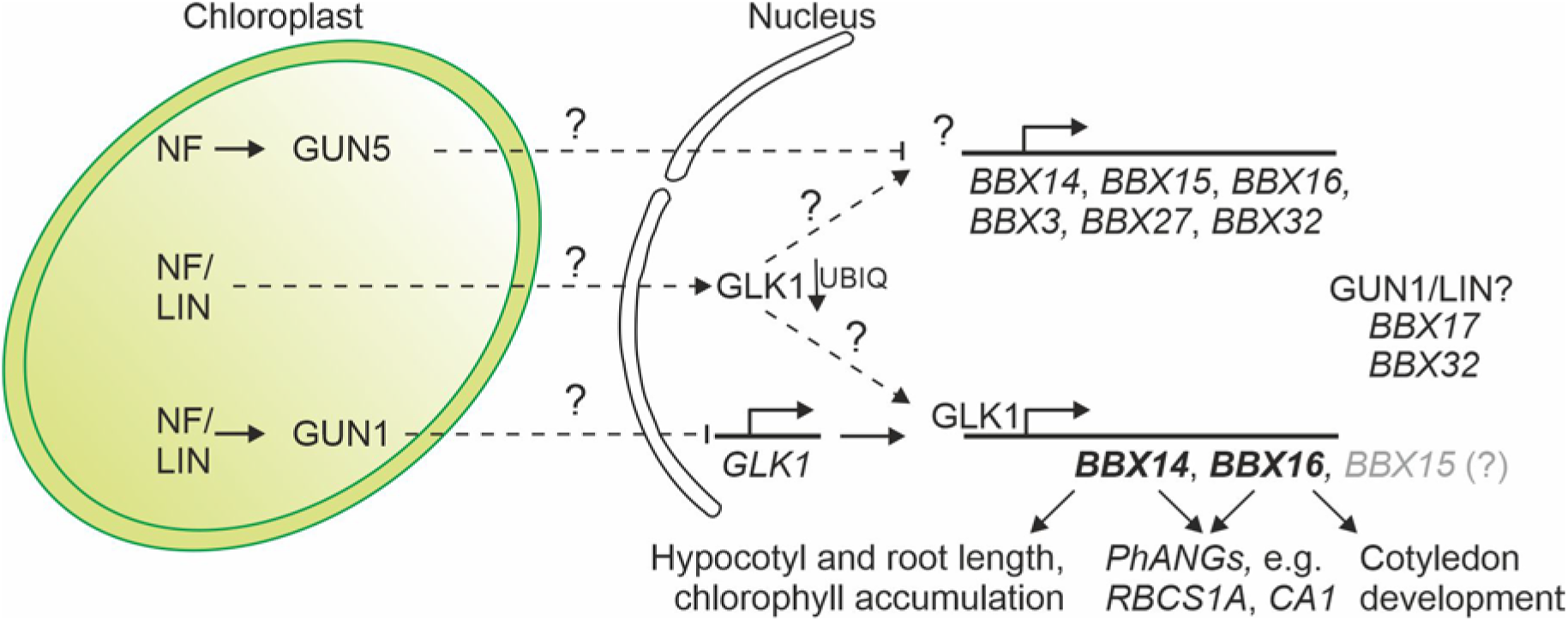
Tentative model for the involvement of BBX proteins in retrograde signaling and seedling photomorphogenesis. Only those *BBX* genes whose mRNA levels are reduced in the presence of norflurazon (NF) only, or in response to NF or lincomycin (LIN) are depicted. As yet unknown signaling pathways are indicated by question marks. The GUN1- and GUN5-dependent pathways are thought to be distinct (Mochizuki *et al*., 2001). While *BBX3*, *BBX14*-*BBX16*, *BBX27* and *BBX32* transcripts are reduced under NF treatment in a GUN5-dependent manner (Fig. S6), *BBX17* is reduced solely by LIN. *BBX14*-*BBX17* all respond to either NF or LIN, and the reduction of *BBX14*-*BBX17* expression is GUN1-dependent, at least under NF conditions. In our ChIP-Seq set-up only *BBX14*-*BB16* were targeted by GLK1. Therefore, it is uncertain whether BBX17 and BBX32 also participate in the GUN1/GLK1 module. Overexpression of BBX14 or BBX16 results in partial *gun* phenotypes by de-repressing photosynthesis-associated nuclear genes (*PhANG*s), while this function remains to be determined for BBX15. Dissection of the retrograde signaling pathway may be hampered by complex interplays and feedback loops. GLK1 transcript and protein levels are reduced for example in a GUN1-dependent and -independent manner, respectively (see Discussion).

Overall, a picture is emerging in which BBX proteins have important roles in plant responses to stresses including the blockage of chloroplast development, in addition to their functions in plant development, but our understanding of their specific functions is far from complete. Functional redundancies and feed-back loops complicate their analysis, and further research will be required to reveal the precise regulatory networks and molecular mechanisms that mediate the signaling and its coordinated responses in nuclear gene regulation.

## ACKNOWLEDGEMENTS

Funding was provided by the Deutsche Forschungsgemeinschaft to K.K., D.L., and T.K. (TRR175, projects C01 and C05).

We thank Paul Hardy for critical comments on the manuscript, and Ramona Kandler for excellent technical assistance.

## DATA AVAILABILITY

Sequencing data have been deposited in NCBI’s Gene Expression Omnibus (Edgar *et al*., 2002) and are accessible under the GEO Series accession number <u>GSE225039</u>. Reads from experiments conducted by (Habermann *et al*., 2020), (Richter *et al*., 2020) and (Xu *et al*., 2020) were retrieved from the NCBI Sequence Read Archive (number PRJNA557616) and Gene Expression Omnibus (numbers GSE104868 and GSE130337, respectively).

## AUTHOR CONTRIBUTIONS

Conceptualization, formal analysis and supervision: T.K.; investigation: V.A., J.S., J.M.M., and T.K.; generation of CRISPR lines, L.W.; help with HL experiments, C.L.; writing – original draft: T.K. with input from K.K. and V.A.; – review and editing: D.L. and T.K.; funding acquisition: D.L., K.K. and T.K.

## Supporting Information

**Fig. S1.**
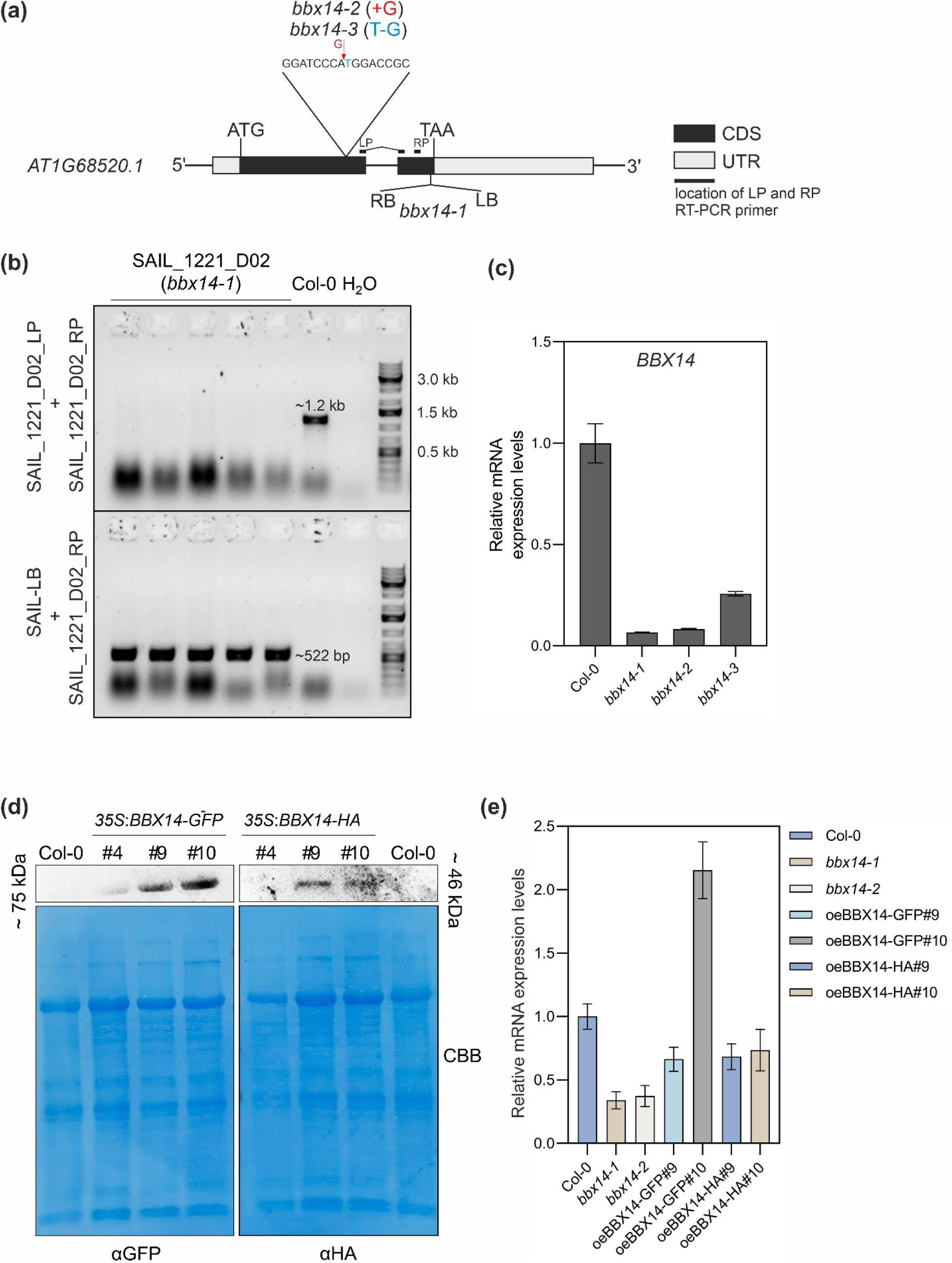
Identification of *bbx14* mutant and *BBX14* overexpression lines. **(a)** Schematic representation of the *BBX14 (AT1G68520)* locus. Exons (black boxes), introns (black lines) and the 5’ and 3’ UTRs (grey boxes) are shown. The location and orientation of the T-DNA insertion in *bbx14-1* are indicated. Note that the insertions are not drawn to scale. Furthermore, the locations of the primers used in the RT-qPCR analysis of *BBX14* expression shown in panels (c) and (e) are indicated as thick black lines. The LP primer is an exon-exon primer spanning intron 1 of *BBX14*, which avoids amplification of genomic DNA. The target sequence of the gRNA used to generate CRISPR/Cas mutant lines is also indicated. The context of the nucleotide insertion in *bbx14-2* (red arrow), as well as the nucleotide substitution T/G in *bbx14-3* (blue) as determined by Sanger sequencing, is shown. **(b)** Confirmation and identification of the homozygous T-DNA insertion in the *bbx14-1* T-DNA line. The gene-specific left and right primers (SAIL_1221_D02_LP and _RP) were used for PCR amplification of sequences around the T-DNA insertion. RP was used together with the T-DNA left border primer (SAIL-LB) for sequencing of the PCR product and verification of the T-DNA insertion. **(c)** RT-qPCR of *BBX14* mRNA expression in 7-day-old wild-type Col-0 and *bbx14* mutant seedlings grown under control conditions (16-h light/8-h dark, 100 µmol photons m^−2^ s^−1^). The results were normalized to *AT4G36800*, which encodes a RUB1-conjugating enzyme (RCE1). Expression values are reported relative to the corresponding transcript levels in Col-0, which were set to 1. Mean values ± SE were derived from two independent experiments (n=2), each performed with three technical replicates per sample. Statistically significant differences (Tukey’s test; \*\**P* < 0.01, \**P* < 0.05) between Col-0 and mutant samples are indicated. **(d)** Total leaf proteins were isolated from 2-week-old Col-0 and Col-0 plants transformed with constructs containing *BBX14-eGFP* or *BBX14-HA* fusions, which were placed under the control of the *35S* promoter. Aliquots were fractionated by 10% SDS-PAGE under reducing conditions, and subjected to immunoblotting using antibodies raised against the GFP- or HA-tag, respectively. PVDF membranes were stained with Coomassie brilliant blue (CBB) to control for protein loading. Representative blots from two experiments are presented. Relative sizes of the BBX14-GFP and BBX14-HA fusion proteins are indicated. **(e)** RT-qPCR of *BBX14* mRNA expression in 7-day-old wild-type (Col-0), *bbx14-1*, *bbx14-2*, and in Col-0 plants “overexpressing” BBX14 (*oeBBX14*). RT-qPCR was performed as described in **(c)**.

**Fig. S2.**
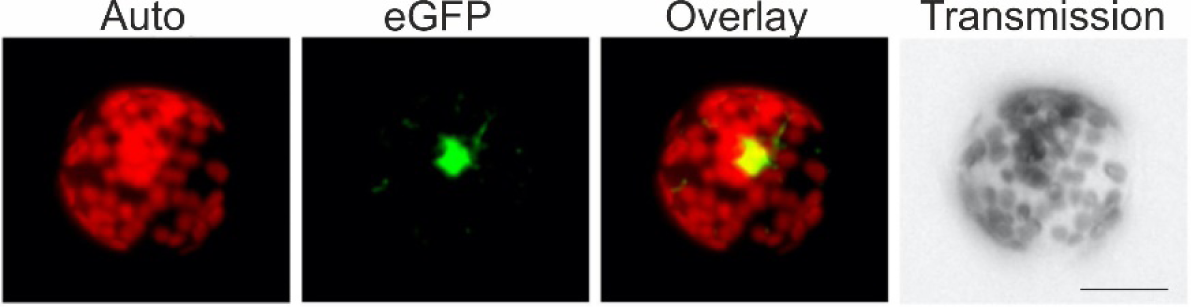
BBX14 is localized to the nucleus. Fluorescence microscopy of tobacco protoplasts transiently expressing BBX14 fused to eGFP. The eGFP fluorescence (green) and chloroplast autofluorescence (red) are shown together in the overlay picture. Scale bar = 10 μm.

**Fig. S3.**
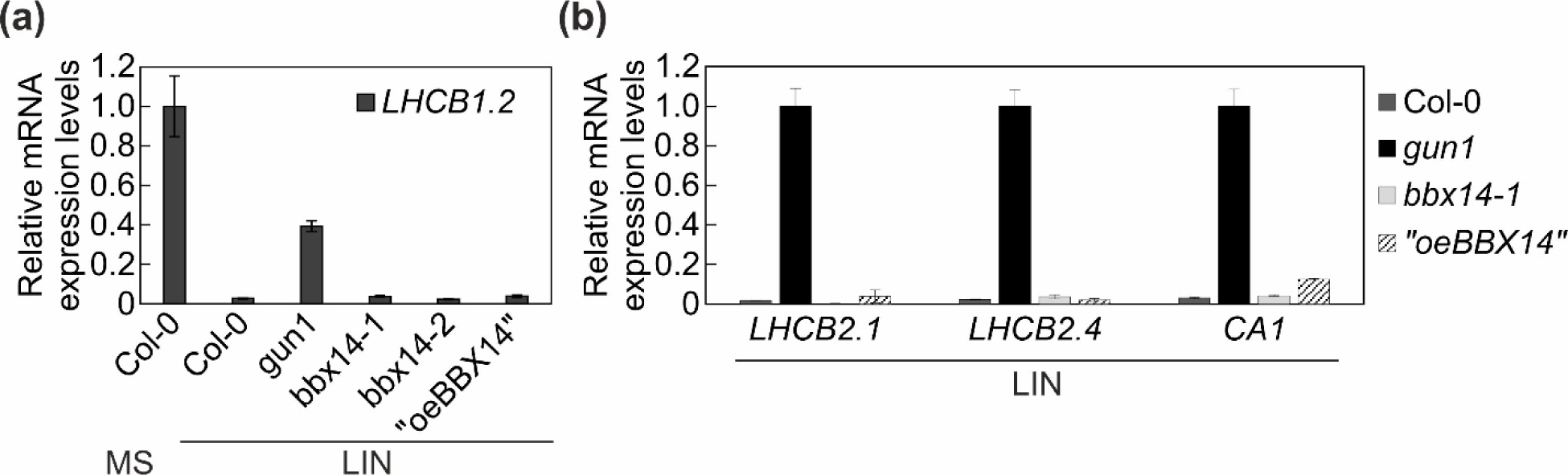
Investigation of putative *gun* phenotypes of seedlings containing altered levels of *BBX14*. RT-qPCR of retrograde marker genes *LHCB1.2* **(a)**, and *LHCB2.2*, *LHCB2.4* and *CA1* **(b)** in Col-0, *gun1* and *bbx14-1* mutants, and the *35S*:*BBX14* line with strongest overexpression (2.5-fold) of *BBX14* (“oeBBX14”). Seedlings were grown for 4 days in continuous light (100 µmol photons m^−2^ s^−1^) in the absence (MS) or presence of lincomycin (LIN). RT-qPCR was performed as described in the legend to Fig. S1c.

**Fig. S4.**
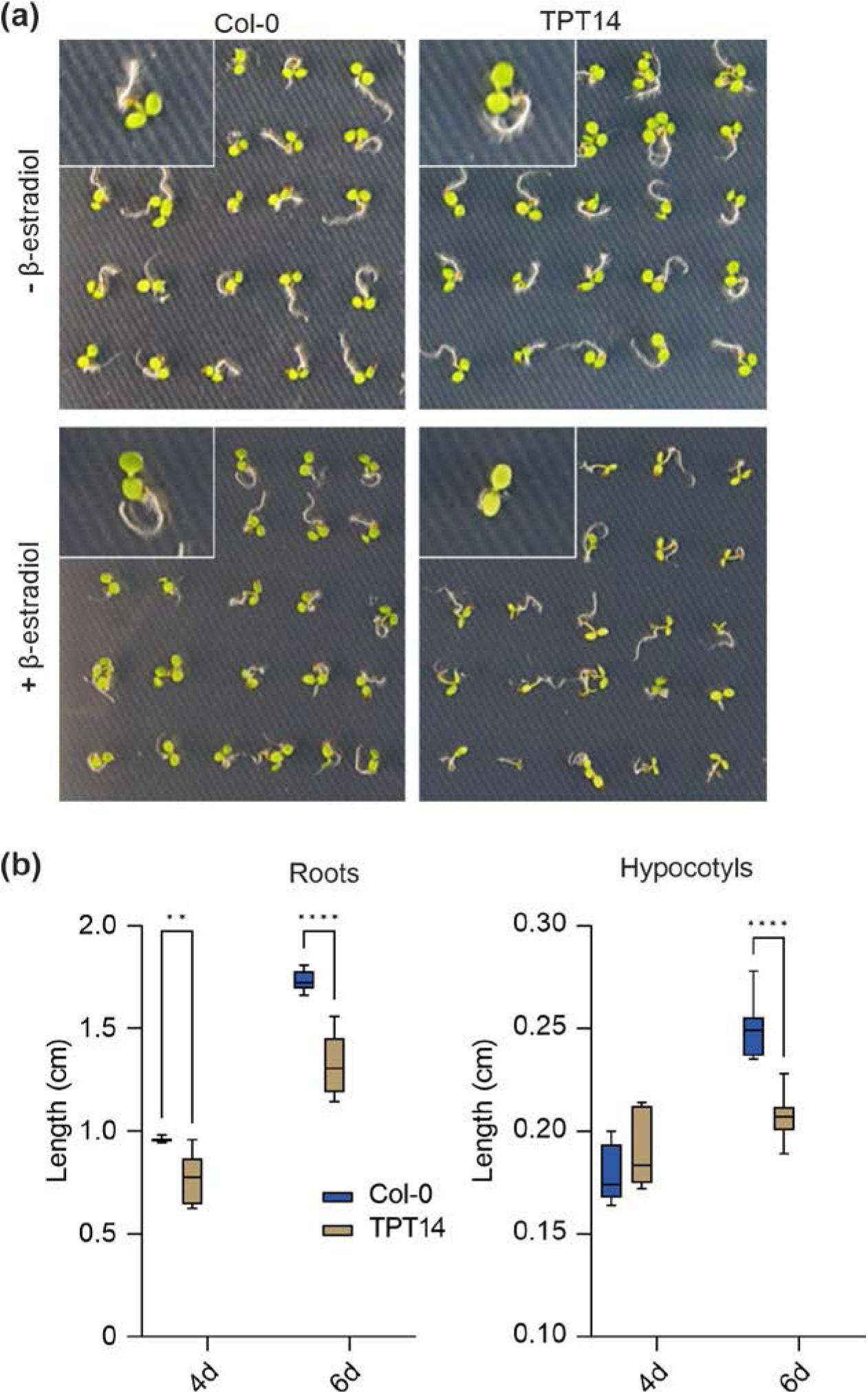
Behavior of the inducible *BBX14* overexpression line. **(a)** Phenotypes of 4-day-old Col-0 and the inducible *BBX14* (TPT14) overexpression line grown under standard conditions (16-h light/8-h dark and 100 µmol photons m^−2^ s^−1^) in the absence (top panel) or presence (bottom panel) of β-estradiol. Scale bars = 0.5 cm. **(b)** Quantification of root and hypocotyl lengths of seedling grown as in **(a)**. The center line of boxplots indicates the median, the box defines the interquartile range, and the whiskers indicate minimum and maximum values. Statistically significant differences between wild-type and the mutant line are indicated (two-way ANOVA; \*\*\*\**P* < 0.0001, \*\*\**P* < 0.001, \*\**P* < 0.01). Scale bar = 0.5 cm.

**Fig. S5.**
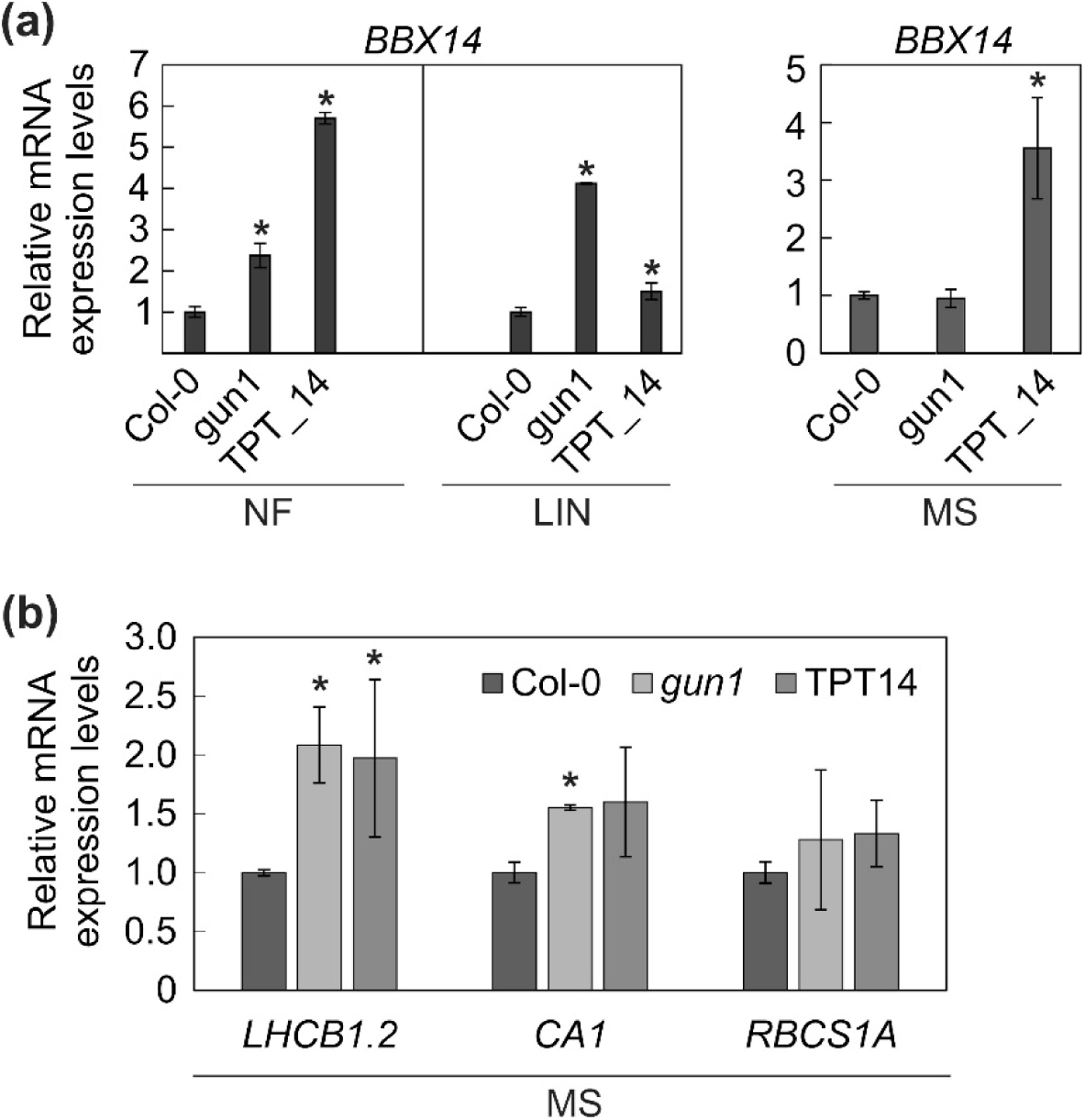
RT-qPCR analysis of **(a)** *BBX14* transcripts in Col-0, *gun1* and *TPT14* lines grown for three days in darkness in the absence of inhibitor, sprayed with the appropriate inducer, put back in the dark for 2 h, and exposed to light (100 µmol m^-2^ s^-1^) for 16 h, and **(b)** *RBCS1A* and *CA1* transcripts of seedlings treated as in (a), but grown on control conditions. The results were normalized to *RCE1*. Expression values are reported relative to the corresponding transcript levels in Col-0, which were set to 1. Mean values were derived from two to three independent experiments, each with three technical replicates. Bars indicate standard deviations. Statistically significant differences (Tukey’s test; *P* < 0.05) between Col-0 and mutant samples are indicated by an asterisk.

**Fig. S6.**
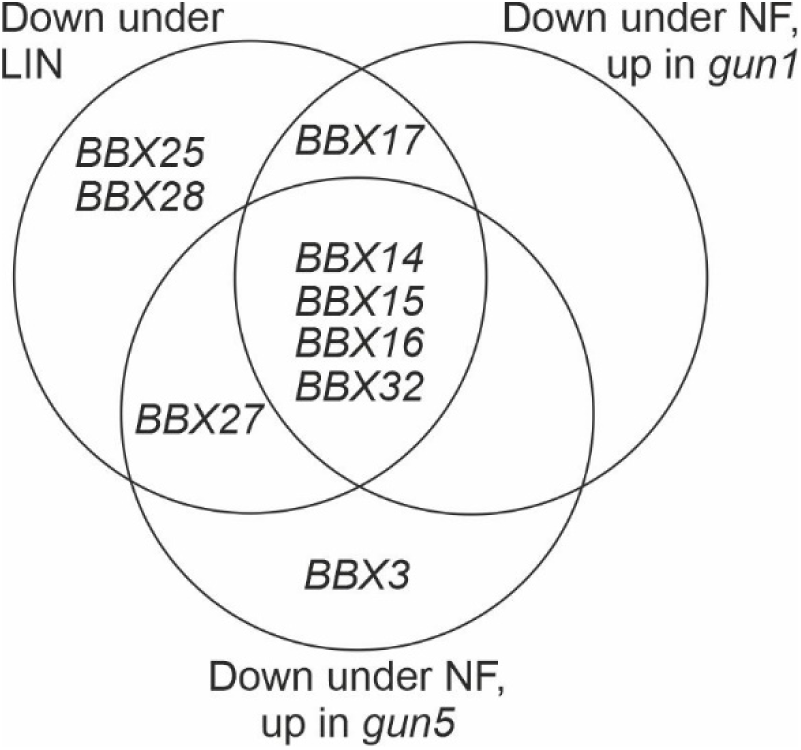
Expression behavior of other *BBX* members under NF and LIN conditions. Venn diagram depicting the degree of overlap between the sets of *BBX* genes whose expression levels were at least two-fold reduced under lincomycin treatment (down under LIN) or at least two-fold reduced under norflurazon (NF), but 2-fold up-regulated when *gun1* or *gun5* were compared to Col-0, respectively (down under NF, up in *gun1* or down under NF, up in *gun5*, respectively). Reads from experiments conducted by (Habermann *et al*., 2020), (Richter *et al*., 2020) and (Xu *et al*., 2020) were retrieved from the NCBI Sequence Read Archive (number PRJNA557616) and Gene Expression Omnibus (numbers GSE104868 and GSE130337, respectively).

**Table S1.**
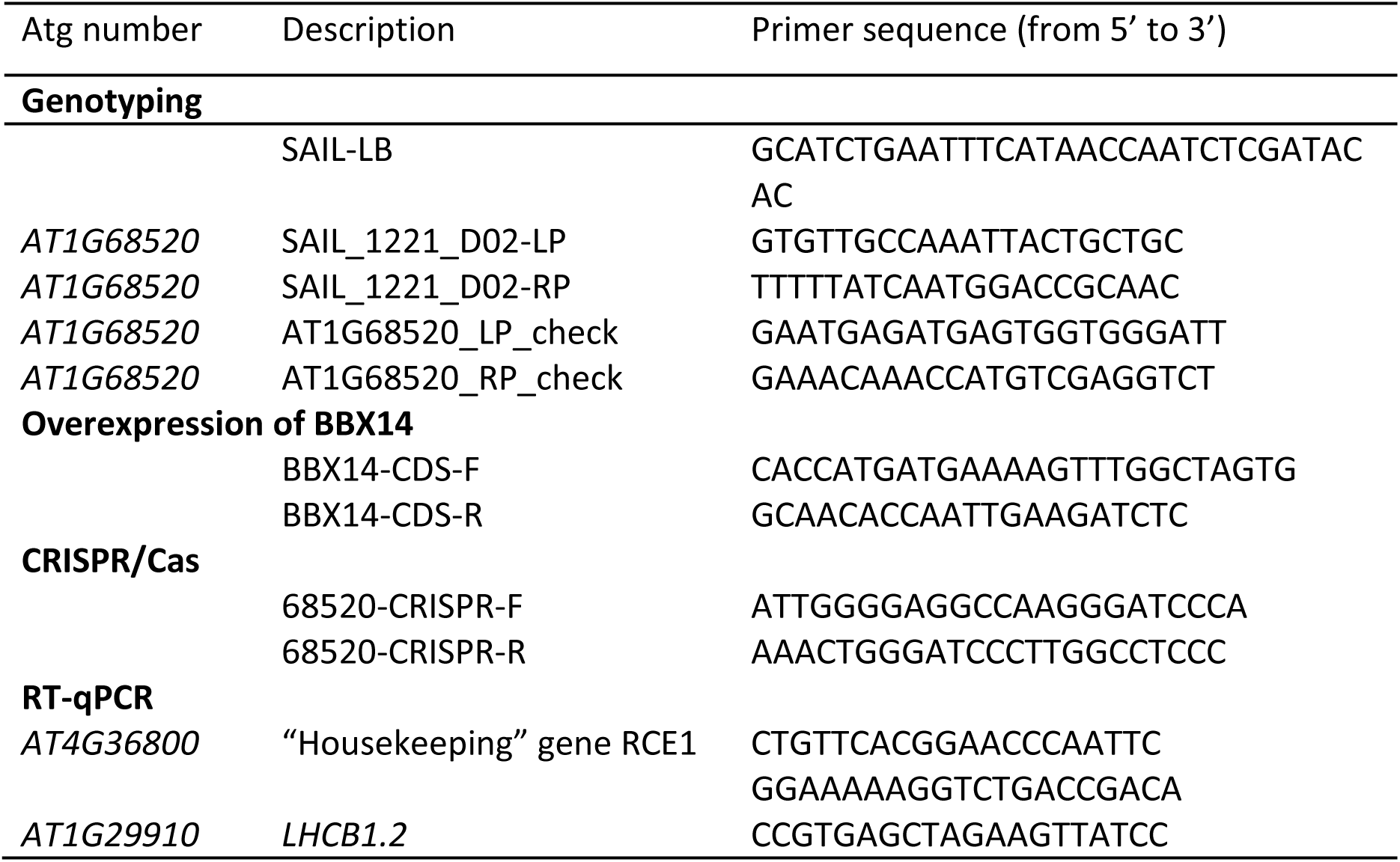
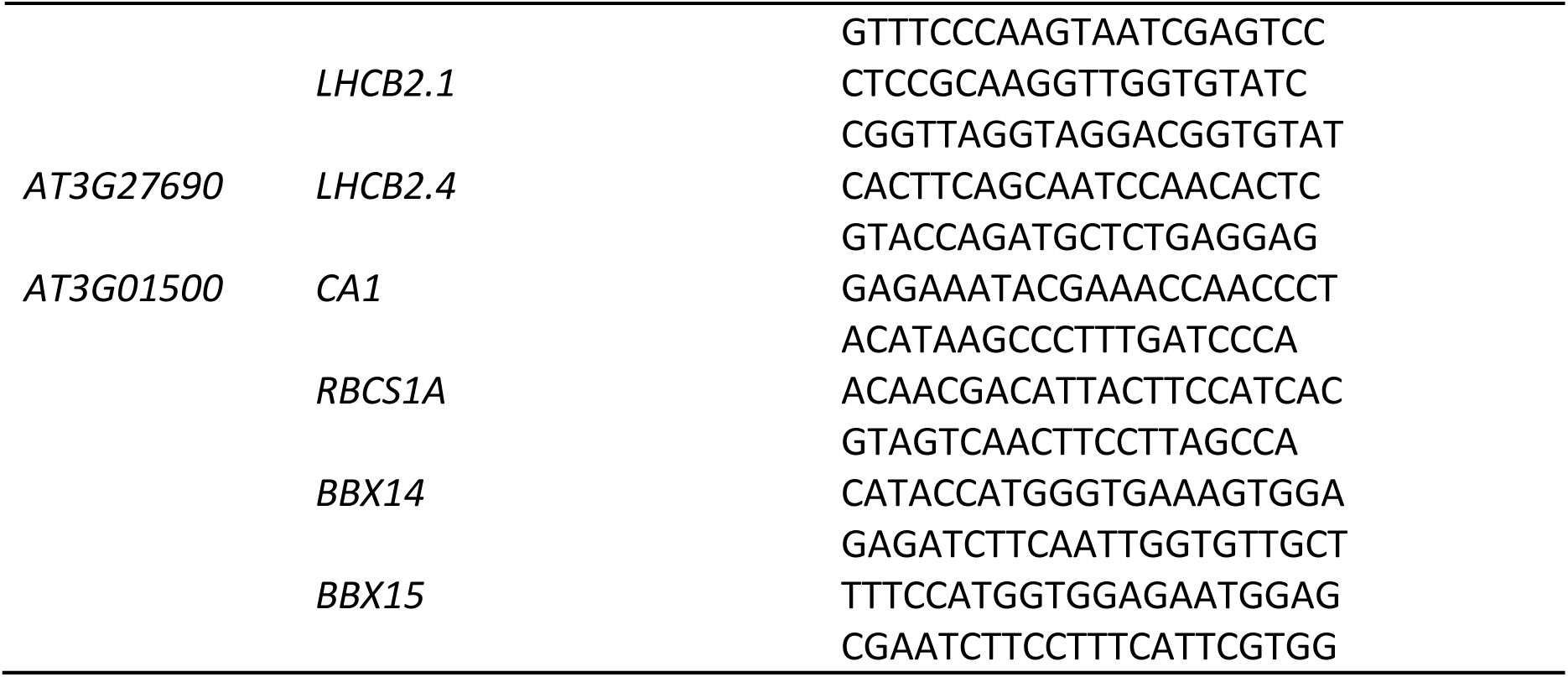
Primers used in this study.

**Table S2** GLK1 ChIP-seq data analysis. Peaks with adjusted *P* values < 0.05 are shown. Analysis was performed by DESeq 2 of both ChIP-seq samples vs. input DNA.

**Table S3** Genes whose transcript levels were significantly changed in 3-day-old etiolated *bbx14-1* seedlings compared to Col-0.

**Table S4** Genes whose transcript levels were significantly changed in 3-day-old etiolated *glk1 glk2* seedlings compared to Col-0.

**Table S5** Genes whose transcript levels were significantly changed in 3-day-old etiolated *bbx14-1* and *glk1 glk2* seedlings that were shifted into the light for 1 day.

## Methods S1 Plant growth conditions

Seeds were surface-sterilized in 10 % bleach and 0.01% Triton-X-100 for 10 min, they were plated on sucrose-free agar (Sigma-Aldrich/Merck, Darmstadt, Germany) plates containing 0.5X Murashige and Skoog (MS) medium followed by stratification at 4°C in dark for 3 days and growth at 22°C under 100 µmol photons m^-2^ sec^-1^ provided by white fluorescent lamps. In the case of dark treatment, seeds were exposed to 2 h of white light (100 µmol photons m^-2^ sec^-1^) after stratification, before allowing germination in the dark. For experiments involving lincomycin or norflurazon treatment, medium was supplemented with 0.5 mM lincomycin (Sigma-Aldrich) or 5µM norflurazon (Sigma-Aldrich). Temperature (22°C/20°C during the day/night cycle) and relative humidity (60%) were strictly controlled under all conditions. For high light experiments, plants were pre-grown for 1 week in a LED chamber set to 80 µmol photons m^-2^ s^-1^; 16 h light/8 h dark cycle), and then irradiance was increased to 1000 µmol photons m^-2^ s^-1^ under strictly controlled temperature conditions.

In experiments performed with inducible overexpression lines, overexpression was induced by supplementing 0.5X MS medium with 2.5 µM β-estradiol or by spraying 4-d-old seedlings with a solution containing 20 µM β-estradiol (Sigma-Aldrich), 0.01 % Silwet L-77 (Lehle seeds) and 0.2% DMSO. In case of experiments shown in Fig. 8, induction was performed immediately after 3 days of dark treatment, followed by 2 h of dark incubation and growth for ≥ 16 hours at 100 µmol photons m^-2^ sec^-1^ continuous white light.

## Methods 2 Protein isolation and immunoblot analyses

Proteins were homogenized in 2× Laemmli sample buffer (120 mM Tris-HCl, pH 6.8, 4% SDS, 20% glycerol, 2.5% ß-mercaptoethanol, 0.01% bromophenol blue), incubated for 10 min at 95°C and centrifuged for 10 min. Proteins were fractionated in a 10% (w/v) SDS-polyacrylamide gel and transferred to PVDF (polyvinylidene fluoride) membranes (Millipore, Billerica, Mass., USA) via semi-dry western transfer using Trans-Blot Turbo (Bio-Rad) in a buffer containing 25 mM Tris, 190 mM glycine, 0.1% and 20% methanol. Membranes were blocked with 5% (w/v) milk in TBS-T (10 mM Tris, pH 8.0, 150 mM NaCl, and 0.1% Tween 20), and probed with monoclonal anti-HA (G1546; Sigma-Aldrich) and anti-GFP (A6455; Life Technologies) antibodies in 1:1000 and 1:5000 dilutions, respectively. Protein loading and transfer were verified by staining PVDF membranes with Coomassie brilliant blue R-250 dye. Signals were visualized via enhanced chemiluminescence Pierce™ ECL Western-Blotting substrate reagent (ThermoFisher Scientific, Waltham, MA, USA) using an ECL reader system (Fusion FX7, PeqLab). Signals were and quantified using ImageJ software (http://rsbweb.nih.gov/ij).

## Methods S3 ChIP-seq sample preparation and data analysis

Seedlings (pGLK1:gGLK1-GFP in Col-0) were grown for 14 days under LD conditions (16 h light, 8 h dark) at 22 °C. To assess whether different sugar and proteasome inhibitor treatments have an effect on GLK1-GFP stability and thus on the number and positions of identified DNA-binding sites, two growth conditions were applied: Half of the seedlings were grown on solid 0.5x MS medium supplemented with 3% sucrose until harvest at ZT8 on day 14 (“untreated”). The second half of the seedlings were grown on solid 0.5x MS medium without sucrose. On day 14 at ZT2, these seedlings were transferred to liquid 0.5 MS medium supplemented with 5 % sucrose and 50 µM MG-132. They were then incubated for another 6 hours in light until harvest at ZT8 (“treated”). 2.5-3 g of MG-132 treated vs. untreated seedlings were harvested on ice and fixed as described (Kaufmann *et al*., 2010). Chromatin immunoprecipitation experiments and library preparation were performed using a GFP antibody (abcam 290) as described (Yan *et al*., 2018). Input DNA was processed in parallel for library preparation.

Fastq files were trimmed from adapter sequences using *Trimmomatic* v0.39 (Bolger *et al*., 2014). Reads were mapped to the Arabidopsis genome (TAIR10) using Bowtie2 v2.2.5 (Langmead & Salzberg, 2012) with default parameters. Only reads mapping to the nuclear chromosomes and with a mapping quality (MAPQ) > 40 were considered for further analysis. Peak calling was done with MACS2 v2.2.7.1 (Zhang *et al*., 2008) comparing IP versus control samples for each tissue independently with parameters: *-p=0.05, -c=220527*. Later, all the identified regions were merged in one file using *mergeBed* from the *BedTools* package (Quinlan & Hall, 2010). *FeatureCounts* (Liao et al., 2014) with parameters *-s 0* was used to obtain the number of mapped reads per region and ChIP-seq and control sample. *DEseq2* (Love *et al*., 2014) was used to detect regions with different number of reads between the IP samples and control. For each binding site, genes around the 1-kb or 3-kb binding regions are reported.

